# Local adaptation in populations of *Mycobacterium tuberculosis* endemic to the Indian Ocean Rim

**DOI:** 10.1101/2020.10.20.346866

**Authors:** Fabrizio Menardo, Liliana K. Rutaihwa, Michaela Zwyer, Sonia Borrell, Iñaki Comas, Emilyn Costa Conceição, Mireia Coscolla, Helen Cox, Moses Joloba, Horng-Yunn Dou, Julia Feldmann, Lukas Fenner, Janet Fyfe, Qian Gao, Darío García de Viedma, Alberto L. Garcia-Basteiro, Sebastian M. Gygli, Jerry Hella, Hellen Hiza, Levan Jugheli, Lujeko Kamwela, Midori Kato-Maeda, Qingyun Liu, Serej D. Ley, Chloe Loiseau, Surakameth Mahasirimongkol, Bijaya Malla, Prasit Palittapongarnpim, Niaina Rakotosamimanana, Voahangy Rasolofo, Miriam Reinhard, Klaus Reither, Mohamed Sasamalo, Rafael Silva Duarte, Christophe Sola, Philip Suffys, Karla Valeria Batista Lima, Dorothy Yeboah-Manu, Christian Beisel, Daniela Brites, Sebastien Gagneux

## Abstract

Lineage 1 (L1) and 3 (L3) are two lineages of the *Mycobacterium tuberculosis* complex (MTBC), causing tuberculosis (TB) in humans. L1 and L3 are endemic to the Rim of the Indian Ocean, the region that accounts for most of the world’s new TB cases. Despite their relevance for this region, L1 and L3 remain understudied. Here we analyzed 2,938 L1 and 2,030 L3 whole genome sequences originating from 69 countries. We show that South Asia played a central role in the dispersion of these two lineages to neighboring regions. Moreover, we found that L1 exhibits signatures of local adaptation at the *esxH* locus, a gene coding for a secreted effector that targets the human endosomal sorting complex, and is included in several vaccine candidates. Our study highlights the importance of genetic diversity in the MTBC, and sheds new light on two of the most important MTBC lineages affecting humans.

## Main

The global tuberculosis (TB) epidemic is a major public health emergency, disproportionately affecting vulnerable populations and worsening existing inequalities. Although the estimated incidence and the number of fatalities have slowly decreased over time, every year ten million people develop the disease, and 1.4 million TB patients die (WHO 2020). Moreover, the Covid-19 pandemic will likely set back the progress toward TB eradication for several year (Glaziou 2020, Hogan et al. 2020, McQuaid et al. 2020, Saunders et al. 2020). TB is spread worldwide, but not all regions are equally affected: 95% of new TB cases occur in Africa and Asia, and the eight countries with the largest burden account for two thirds of the total number of cases (WHO 2020).

Human TB is caused by members of the *Mycobacterium tuberculosis* complex (MTBC), which includes nine phylogenetic lineages with different geographic distributions. Five of these lineages are restricted to Africa (L5, L6, L7, L8, and L9), while the remaining four (L1, L2, L3, L4) are more broadly distributed (Chihota et al. 2018, Gagneux 2018, Devis et al. 2020, Ngabonziza et al. 2020). While L2 and L4 strains occur across the world, L1 strains are predominantly found around the rim of the Indian Ocean (East Africa, South Asia, and Southeast Asia). L3 has a distinct geographic range (East Africa, Central Asia, Western Asia, and South Asia) that overlaps partially with L1 (Wiens et al. 2018, Couvin et al. 2019). While many MTBC lineages also occur in Northern Europe, North America, Australia and New Zealand, in these low-burden regions, the majority of TB cases are imported through recent migrations, and local transmission is rare (White et al. 2017).

Beyond their geographic distribution, MTBC lineages also differ in virulence, transmissibility, association with drug resistance, and the host immune responses they elicit (Peters et al. 2020). There is an increasing notion that MTBC genetic variation should be considered in the development of new antibiotics and vaccines, and when studying the epidemiology and pathogenesis of the disease (Gagneux 2017, Wiens et al. 2018, Peters et al. 2020). Yet, much of TB research to date has focused on the laboratory strain H37Rv (belonging to L4) and a few other strains belonging to L2 and L4, because of their broad geographic range, and their association with drug resistance. By contrast, L1 and L3 have largely been neglected, and only few studies have investigated the global populations of these two lineages (Couvin et al. 2019, O’Neill et al. 2019). This knowledge gap is particularly severe as L1 and L3 are endemic to some of the world regions with the heaviest TB burden. For example, L1 and L3 cause the majority of TB cases in India, the country with the highest number of new TB cases in the world, and L1 is by far the most important cause of TB in the Philippines, the country with the 4^th^ highest global TB burden (Wiens et al. 2018). The aim of this study was to characterize the global population structure of L1 and L3 using large-scale population genomics, and to investigate the evolutionary history and selective forces acting on these two lineages.

## Results

We screened a large collection of publicly available MTBC genome sequences and selected those belonging to L1 and L3. Additionally, we newly sequenced 767 strains to cover regions that were not well represented in the public data. We applied a series of bioinformatic filters to exclude low quality sequences and mixed infections (Methods), and obtained a curated genome dataset of 4,968 strains (2,938 L1, 2,030 L3). While the phylogenetic tree of L3 showed a ladder-like topology without an evident division in sublineages, L1 comprised five clearly distinct sublineages coalescing close to the root of the tree (Sup. Figs. 1-2).

The genome sequences included in our final dataset represented 69 countries, and covered the known geographic range of L1 and L3 (Methods, Sup. Table 1, Sup. Figs. 3-4). Because we were interested in studying L1 and L3 in their endemic range, we excluded strains originating from North America, Europe, and Australia from the biogeographic analysis, as in these low-burden regions, most TB patients are recent migrants who were infected in their country of origin (White et al. 2017). We assigned strains to different geographic regions following a modified version of the United Nation geographic scheme (Methods, Figs. 1 and 2). Mapping the regions of origin onto the phylogenetic trees revealed that two sublineages of L1 are found almost exclusively in Southeast Asia (L1.1.1 and L1.2.1), while the others are spread over the complete geographic range of L1 (Fig. 1, Sup. Figs. 5-6).

**Figure 1.**
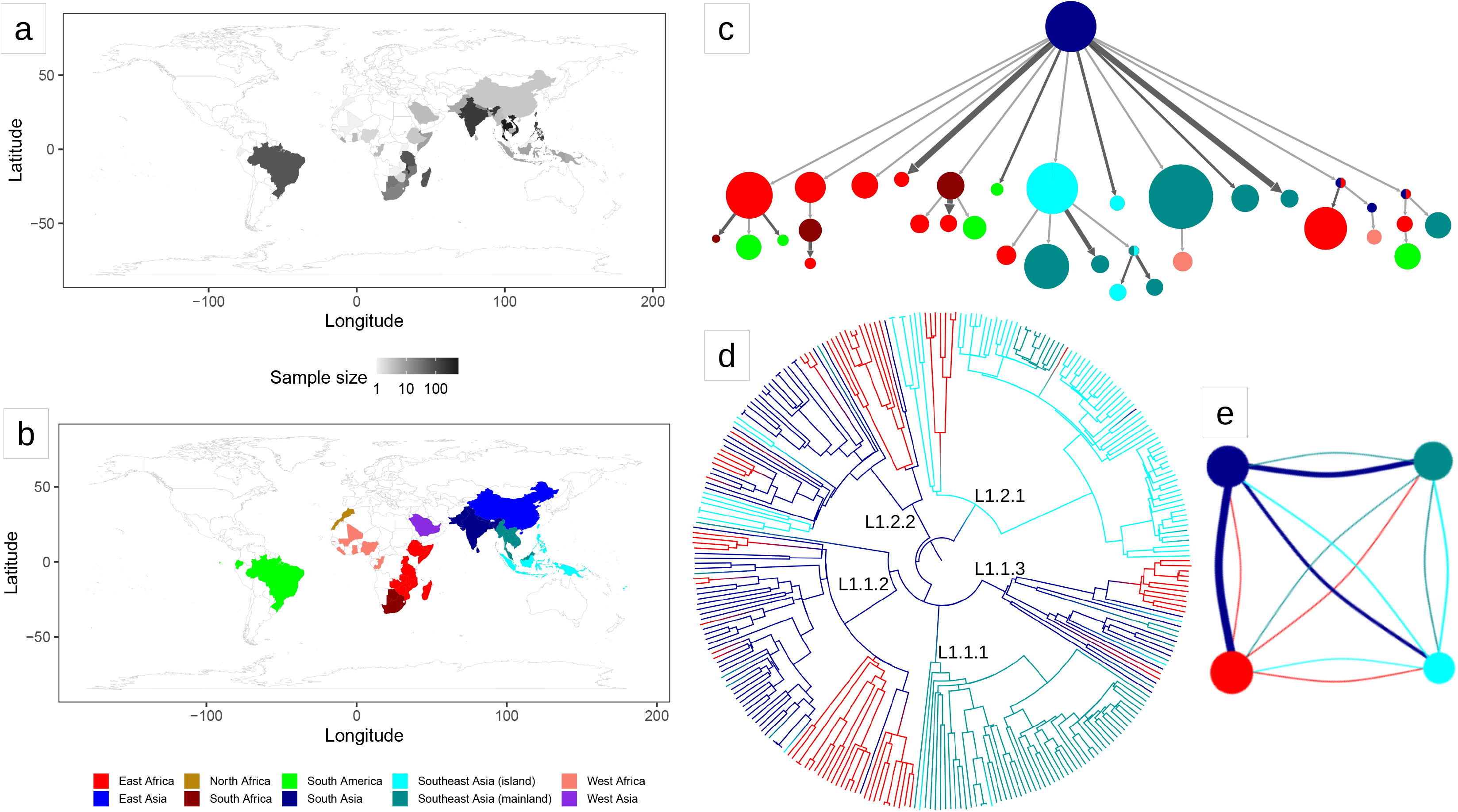
Results of the biogeography analysis: L1. **a)** Heatmap indicating the origin of the 2,061 L1 strains included in the dataset used for the biogeography analysis. **b)** The geographic regions used in the biogeography analysis of L1. **c)** Results of PASTML (compressed tree), the color code follow the legend of panel (b), the size of the circles is proportional to the number of tips, and the size of the arrows is proportional to the number of times the pattern of migration was observed on the tree. This plot excluded nodes that represented less than three strains. The same plot including all nodes can be found as Sup. File 1. **d)** Tree obtained with the Mascot analysis; the branches are colored following the legend in panel (b) and represent the inferred ancestral region with the largest posterior probability. **e)** Representation of the relative effective population sizes (circles) and migration rates (lines connecting circles) estimated by Mascot. Lines representing migration rates are colored based on the region of origin (interpreted forward in time). For example, the dark blue line connecting the dark blue circle with the red circle represents the forward migration rate between South Asia and East Africa, which is the largest migration rate estimated by Mascot. The parameter estimates are reported in Sup. Table 2.

**Figure 2.**
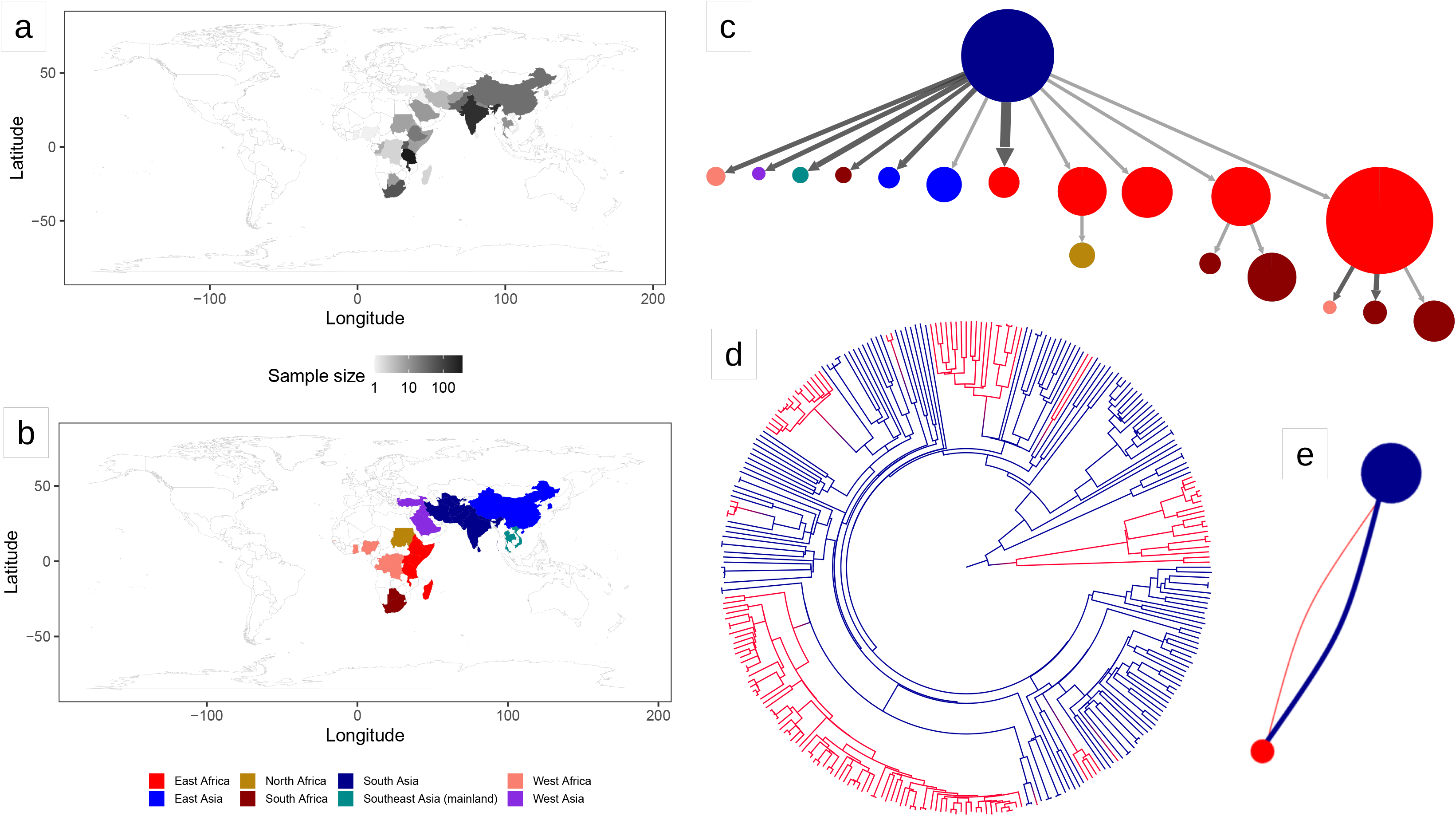
Results of the biogeography analysis: L3. **a)** Heatmap indicating the origin of 1,021 L3 strains included in the dataset used for the biogeography analysis. **b)** The geographic regions used in the biogeography analysis of L3. **c)** Results of PASTML (compressed tree), the color code follow the legend of panel (b), the size of the circles is proportional to the number of tips, and the size of the arrows is proportional to the number of times the pattern of migration was observed on the tree. This plot excluded nodes that represent less than three strains, the same plot including all nodes can be found as Sup. File 3. **d)** Tree obtained with the Mascot analysis; the branches are colored following the legend in panel (b) and represent the inferred ancestral region with the largest posterior probability. **e)** Representation of the relative effective population sizes (circles) and migration rates (lines connecting circles) estimated by Mascot. Lines representing migration rates are colored based on the region of origin (interpreted forward in time). The effective population size of South Asia was estimated to be much larger than the one of East Africa. Additionally, the forward migration rate between South Asia and East Africa was much larger than the one in the opposite direction. The parameter estimates are reported in Sup. Table 2.

We performed a phylogeographic analysis using two alternative approaches (Mascot and PASTML), which are based on different models and inference methods (Müller et al. 2018, Ishikawa et al. 2019). We found that South Asia was predicted to be the geographic range of the Most Recent Common Ancestor (MRCA) of both L1 and L3 (Sup. Information). Most interestingly, we found a strongly asymmetric pattern of migration. For L1, PASTML identified most migration events from South Asia toward other regions, followed by further dispersion, but almost no back migration to South Asia (Fig. 1; Sup. Fig. 7; Sup. Files 1-2). When we estimated the migration rates with Mascot, we found that the forward migration rate from South Asia toward the rest of the world were 3 to 17 times larger than the migration rates in the opposite direction, confirming the results of PASTML (Fig. 1, Sup. Table 2). We found a similar scenario for L3: PASTML inferred the largest number of migrations from South Asia toward East Africa, further spread from East Africa toward neighboring regions, but essentially no migration back to South Asia (Fig. 2; Sup. Files 3-4). The Mascot analysis showed that the forward migration rates from South Asia toward East Africa were 26 times larger than the migration rates in the opposite direction (Fig. 2, Sup. Table 2).

We performed tip dating to estimate the age of the trees but got inconsistent results for L1, due to a lack of a reliable temporal signal (Sup. Information, Sup. Figs. 8-9, Sup. Tables 3-4). However, the results of a previous study suggested a relatively fast evolutionary rate for L1 (~ 1.4×10^−7^ nucleotide changes per site per year; Menardo et al. 2019), and that the MRCA of L1 lived around the 12^th^ century AD (Sup. Information). By contrast, L3 had a good temporal signal, and different methods estimated that its MRCA lived between the 2^nd^ and the 13^th^ century AD. However, the uncertainty around all of these estimates was very large (Sup. Information, Sup. Fig. 8, Sup. Tables 5-6). Together, these results corroborate the findings of previous studies (Bos et al. 2014, Duchêne et al. 2016, Menardo et al. 2019). Calibrating MTBC trees that are hundreds or thousands of years old, with sequences sampled in the last few decades is notoriously challenging and subject to limitations (Menardo et al. 2019). Therefore, we refrain from any strong interpretation of the results of the molecular clock analyses of L1 and L3. We emphasize that the ages reported here are the most likely estimates supported by the available data, and additional data, or alternative methods, might result in different temporal scenarios.

With respect to the geographical aspects, we identified several interesting instances of long-range dispersal. First, we found a clade of L1, composed of 11 strains sampled in five different West African countries. This was surprising because West Africa is usually not considered part of the geographic range of L1. This clade is nested within sublineage L1.1.1, which is essentially only found in the mainland of Southeast Asia, and the PASTML analysis inferred a direct introduction from Southeast Asia to West Africa. This introduction is unlikely to have happened before the 16^th^ century, when the Portuguese established the maritime route between Europe and Asia by circumnavigating Africa. Assuming that this migration did not occur before the 16^th^ century, we benchmarked the molecular clock of L1 and found that this scenario is indeed compatible with a clock rate equal to, or larger than 1.4×10^−7^ nucleotide changes per site per year, but not with lower ones (Sup. Information, Sup. Fig. 10). Second, we found that L1 was introduced to South America at least on 11 independent occasions (Fig. 1, Sup. Files 1-2). Assuming a clock rate of 1.4×10^−7^, the earliest introduction was between 1620 and 1830 AD from East Africa, while subsequent introductions occurred from East Africa, South Africa, and South Asia. These results support the hypothesis that L1 was first introduced to Brazil through the slave trade from East Africa (Allen 2013, Conceição et al. 2019). Interestingly, this is in contrast with L5 and L6, which are endemic to West Africa, but did not establish themselves firmly in South America (De Jong et al. 2010, Rabahi et al. 2020).

Third, similar to the West African clade of L1.1.1, we found an East African clade embedded within sublineage L1.2.1, which otherwise is found almost exclusively in Southeast Asia. This East African clade is composed of 11 strains from five countries, and its sister clade is found in East Timor and Papua New Guinea. We inferred a direct migration from the islands of Southeast Asia to East Africa that occurred between the 13^th^ and the 16^th^ century AD (assuming a clock rate of 1.4×10^−7^). This would be compatible with early Portuguese expeditions, which reached East Timor and Papua New Guinea in the early 16^th^ century. An alternative explanation could be the Austronesian expansion. Austronesians are thought to have reached the Comoros islands and Madagascar between the 9^th^ and the 13^th^ century AD, possibly through direct navigation from Southeast Asian islands. Malagasy speak an Austronesian language, and Austronesian genetic signatures are found in in human populations in the Comoros, Madagascar, and to a small extent also in the Horn of Africa (Blench 2010, Boivin et al. 2013, Pierron et al. 2014, Brucato et al. 2016, Brucato et al. 2018, Brucato et al. 2019).

L1 and L3 coexist in many regions around the Indian Ocean. Yet, in their evolutionary history these lineages colonized areas occupied by different human populations. Human genetic variation has been shown to influence the susceptibility to TB (Qu et al. 2011). Most notably, HLA genes play a crucial role in the activation of the immune responses to the MTBC by exposing bacterial peptides (epitopes) to the surface of an infected cell, where they can be recognized by T cells. HLA genes are extremely polymorphic in human populations, and several alleles of different HLA genes are associated with TB susceptibility in different populations (Brahmajothi et al. 1991, Vejbaesya et al. 2002, Yuliwulandari et al. 2010, Salie et al. 2014, Sveinbjornsson et al. 2016).

Previous studies have shown that T cell epitopes are hyper-conserved in the MTBC, suggesting that immune escape does not provide an advantage, and that contrary to other pathogens, the MTBC needs to be recognized by the immune system and to cause disease in order to transmit (Comas et al. 2010, Coscolla et al. 2015).

Our large dataset of L1 and L3 genome sequences from different geographic regions provided an opportunity to scan for lineage- and/or region-specific signatures of selection at T cell epitopes in L1 and L3.

We reconstructed the mutational history of T cell epitopes in L1 and L3, and found that 51% of all epitopes were variable (at the amino acid level) in at least one L1 strain, while only 20% were variable in at least one L3 strain (Sup. Table 7). However, this difference can be explained by the different size and diversity of the two datasets (2,061 genome sequences with 136,023 variable sites for L1; 1,021 genome sequences with 36,316 variable sites for L3). The epitope that accumulated most mutations was located at the N-terminus of *esxH* (*Rv0288*). This peptide is a T cell epitope for both classes, MHCI and MHCII; it is also a B cell epitope, and was previously identified as one of the few T cell epitopes that were not hyper-conserved (Comas et al. 2010). We found 15 derived haplotypes (at the amino acid level), generated by 28 independent replacements at five positions in a peptide of seven amino acids (Figure 3). By contrast, the second most variable epitope accumulated only seven amino acid changes (Sup. Table 7). Interestingly, we did not find this signature in L3, as all strains carried the ancestral haplotype at the N-terminal *esxH* epitope. Moreover, when we extended the analysis to two large genomic datasets of L2 and L4 strains, we found a much weaker signal: while 21% of L1 strains carried a derived haplotype for this epitope, only 1% of L2 and L4 strains, and no L3 strain had a derived haplotype. Despite analyzing datasets with more strains (6,752 and 10,466 for L2 and L4, respectively), and more polymorphic positions (140,309 and 277,648) compared to L1, we found only three amino acid replacements at the N-terminal epitope of *esxH* in L2, and seven in L4 (Figure 3). The most frequently mutated position was the tenth amino acid, where we found 12 independent replacements in L1, and two in L2 and L4 (Fig. 3). The amino acid replacement A10V occurred eight times in parallel in L1, once in L2 and twice in L4. The most abundant derived haplotype was caused by a different amino acid replacement at position ten (A10T), which occurred once in L1 and once in L2 (Fig. 3). Overall, the replacements in all lineages occurred in eight residues in a peptide of 13 amino acids (Fig. 3).

**Figure 3.**
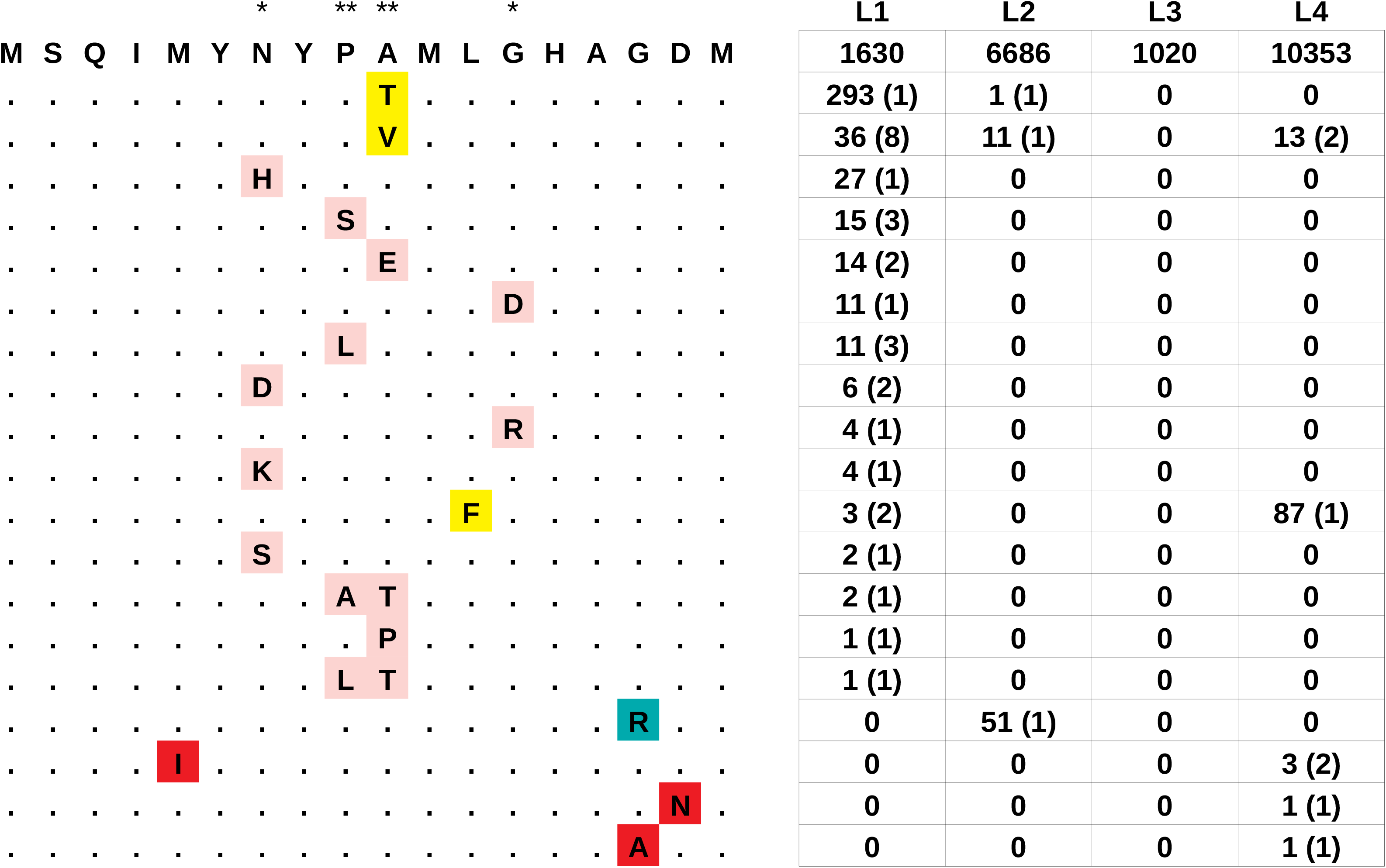
The hypervariable epitope at the N terminus of *esxH* (aa 1-18). The ancestral epitope is reported in top position, the derived haplotypes are reported below: mutations that were present in more than one lineage are highlighted in yellow, haplotypes that were found exclusively in L1, L2 or L4 are highlighted in pink, blue and red, respectively. Asterisks indicate the position inferred to be under positive selection by PAML: * posterior probability > 0.95: ** posterior probability > 0.99. The table on the right reports for each lineage the number of strains harboring the corresponding haplotype, and between parentheses, the number of independent parallel occurrences of the mutations as inferred by PAUP.

We evaluated the robustness of these results by formally testing for positive selection on the complete sequence of *esxH* using PAML (Yang 2007). We found that *esxH* was indeed under positive selection in L1 (p-value = 0.00004) but not in the other lineages (p-values = 0.39, 1.00 and 0.65 for L2, L3 and L4, respectively). PAML identified four codons that have been under positive selection (posterior probability > 0.95), all of them within the N-terminal epitope (codons 7, 9, 10 and 13; Fig. 3). Codon 76, which is part of a different T cell epitope, had a posterior probability > 90%, mutated three times in parallel and was possibly also under positive selection.

Our results further revealed that the derived haplotypes of the N-terminal *esxH* epitope were not distributed randomly across the geographic range of L1. Twenty-two of the 28 (79%) amino acid replacements occurred in sublineages L1.1.1 and L1.2.1, which are almost exclusively present in Southeast Asia (Sup. Fig. 11). We constructed a statistical test of association similar to phyC (Farhat et al. 2013) to determine whether replacements in the hypervariable *esxH* epitope were significantly associated with a particular geographic region (Methods).

We found that South African strains were less likely to harbor a derived haplotype in the N-terminal *esxH* epitope than expected by chance (empirical p-value = 0.0130; Table 1). While East African L1 strains were not associated with the derived haplotypes (empirical p-value = 0.276), we noticed important differences between countries: of the 29 East African strains harboring a derived haplotype, 28 were sampled in Madagascar. When we excluded Madagascar, we found that East Africa had a strong negative association with the derived haplotypes (i.e. East African strains harbored less derived haplotypes than expected by chance; empirical p-value = 0.0004). We then tested the most frequently replaced position (position ten) alone, and again found that East African strains were negatively associated with the derived haplotypes, with and without excluding Malagasy strains (empirical p-values = 0.0176 and 0.034, respectively). Finally, we tested the derived haplotype caused by the most frequent amino acid replacement (A10V; 8 parallel occurrences). Again, we found a negative association with East African strains (empirical p-values = 0.046 and 0.079, respectively including and excluding Malagasy strains; Table 1).

**Table 1.** Results of the test of association between haplotypes of the N-terminal esxH epitope and the geographic region of origin.

| Region<sup>1</sup> | Strains<sup>2</sup> | Association<sup>3</sup> | *esxH* (A10V)<sup>4</sup> |  | *esxH* (A10X)<sup>5</sup> |  | *esxH* (N terminus)<sup>6</sup> |  |
| --- | --- | --- | --- | --- | --- | --- | --- | --- |
|  |  |  | Strains | p-value | Strains | p-value | Strains | p-value |
| South Asia | 219 | + | 1 | 0.4852 | 6 | 0.385 | 17 | 0.453 |
| Southeast Asia (islands) | 235 | + | 10 | 0.154 | 156 | **0.081** | 171 | 0.174 |
| Southeast Asia (mainland) | 982 | + | 25 | 0.200 | 183 | **0.095** | 210 | 0.196 |
| East Africa | 466 | - | 0 | **0.046** | 1 | **0.034** | 29 | 0.276 |
| East Africa (no Madagascar) | 391 | - | 0 | **0.070** | 0 | **0.018** | 1 | **0.0004** |
| South Africa | 49 | - | 0 | 0.305 | 0 | 0.165 | 0 | **0.013** |
| South America | 77 | - | 0 | 0.496 | 0 | 0.352 | 0 | **0.095** |

<sup>1</sup> We included only regions for which the sample size was 25 or more.
<sup>2</sup> Total number of L1 strains from each region.
<sup>3</sup> All p-values are one-tailed empirical p-values. This column indicates the direction of the association: + we tested for positive association between the derived haplotype(s) and the geographic region; - we tested for negative association between derived haplotype(s) and the geographic region.
<sup>4</sup> Number of strains with the derived haplotype caused by the mutation A10V, and results of the test of association. P-values < than 0.1 are shown in bold.
<sup>5</sup> Number of strains with the derived haplotypes caused by any mutation at position 10 (A10X), and results of the test of association. P-values < than 0.1 are shown in bold.
<sup>6</sup> Number of strains with the derived haplotypes caused by any mutations occurred in the first 18 amino acids of EsxH, and results of the test of association. P-values < than 0.1 are shown in bold.

We hypothesized that the accumulation of missense mutations at the N-terminal epitope of *esxH* was due to immune escape. Therefore, we performed *in silico* prediction of the binding affinity of the ancestral haplotype and of the two most frequent derived haplotypes (caused by the amino acid replacement A10V and A10T) with different HLA-A, HLA-B and HLA-DRB1 alleles (Methods). We performed this analysis for:

1. Alleles found at high frequency (> 10%) in South- and Southeast Asia, but not in East Africa.
2. Alleles found at high frequency (> 10%) in East Africa, but not in South- and Southeast Asia.
3. Alleles found at high frequency (> 10%) in both regions. However, we found no differences in the predicted binding affinities between alleles with different geographic distributions (Sup. Table 8).

While *esxH* was the most striking example of a selective pressure specific for one lineage, our analysis suggest that it was not an isolated case. We performed a genome-wide scan for selection with PAML, and identified 17 genes under positive selection, of which five in common between L1 and L3 (Sup. Table 9). We found two genes coding for transmembrane proteins, members of the Esx-1 secretion system, which were under positive selection in L1 (*eccB1* and *eccCa1*, Bonferroni corrected p-values = 0.02 and 0.03), and several genes involved in antibiotic resistance that were under positive selection in both lineages (Sup. Information). We further characterized the profile of drug resistance mutations of L1 and L3, and found that L1 strains harbored a greater proportion of *inhA* promoter mutations (conferring resistance to isoniazid) compared to L3 strains, confirming previous findings (Fenner et al. 2012; Sup. Information; Sup. Fig. 12; Sup. Table 10).

## Discussion

Our results highlight the central role of South Asia in the dispersion of L1 and L3. First, we confirmed that the two lineages probably expanded from South Asia (O’Neill et al. 2019). Second, contrary to previous studies that assumed symmetric migration rates between regions (O’Neill et al. 2019), we found that these migrations occurred mostly in one direction. South Asia was a source of migrant strains that fueled the epidemics in other regions, especially in East Africa. Historically, a network of maritime trade, which followed the seasonality of the Monsoons, connected East Africa and South Asia. It is unclear how this could have promoted the spread of TB in one direction, but not in the other. A possible explanation is that strains originating in South Asia were more efficient in transmitting in East Africa, compared to East African strains that migrated to South Asia.

Another difference compared to previous studies is the temporal framework for the evolutionary history of L1. O’Neill et al. (2019) estimated that the MRCA of L1 lived in the 4^th^ century BC and that the migration rates decreased after the 7^th^ century AD. Our results indicate that the MRCA of L1 did not exist before the 12^th^ century AD. This discrepancy is due to different assumptions about the clock rates of L1, but none of the two hypotheses can be excluded with the available data. Nonetheless, our discovery of a West African clade that originated through a direct introduction from Southeast Asia supports a larger clock rate of L1, and therefore a more recent MRCA (Sup. Information). As we already mentioned, tip-dating analyses of MTBC trees with roots that are several hundreds of years old is extremely challenging, and the results of such analyses should be taken with caution (Menardo et al. 2019). Moreover, all MTBC molecular clock studies so far assumed that evolutionary rate estimates do not depend on the age of the calibration points (Ho et al. 2005). There are some indications that the effect of time dependency in MTBC datasets is negligible (Pepperell et al. 2013, Menardo et al. 2019). However, this assumption has not yet been thoroughly tested due to the lack of appropriate samples.

We found evidence for a strong selective pressure acting on the N-terminus of *esxH* in L1. In contrast, this selective pressure seems to be much reduced, if not absent, in the “modern” lineages (L2, L3 and L4).

It is known that L1 strains interact differently with the infected hosts compared to other lineages. For example, L1 strains show a lower virulence in animal models (Bottai et al. 2020), transmit less efficiently in some clinical settings, and infect older patients (Holt et al. 2018). It has been shown recently that the increased virulence of the so-called “modern” MTBC strains is due to the loss of the genomic region TbD1, which remains present in L1 (Bottai et al. 2020). However, it was also reported that in some populations, L1 was associated with higher patient mortality (Smittipat et al. 2019). Given these differences with the “modern” lineages, it is likely that L1 is subject to different selective pressures, and it is possible that the greater selective pressure on *esxH* was caused by some epistatic effect specific to L1.

The signature of selection at the N-terminal epitope of *esxH* were associated with strains sampled in South- and Southeast Asia, and were almost completely absent in East Africa (excluding Madagascar). Region-specific signatures of positive selection are a hallmark of local adaptation; in this case, adaptation of L1 strains to human hosts with South- and Southeast Asian genotypes. This corroborates previous studies reporting that in Taiwan, L1 is associated with indigenous populations with Austronesian ancestry (reviewed in Dou et al. 2014). This hypothesis is also supported by the L1 population in Madagascar. Madagascar is geographically linked to East Africa, however, Malagasies are genetically distinct from Africans, as they have mixed African and Southeast Asian ancestry due to the Austronesian colonization of Madagascar (Brucato et al. 2016). Madagascar was the region with the second highest prevalence of derived haplotypes in the N-terminal epitope of *esxH* after Southeast Asia (islands), 37.3% and 72.8%, respectively, as opposed to 0.3% (one single strain) in the rest of East Africa.

Although all the codons under positive selection in *esxH* were contained in one single T cell epitope, the selective pressure acting on *esxH* could be due to some other factor, and not to the recognition of the epitope by T cells, for which, indeed, we found no evidence in the binding prediction analysis. EsxH is a small effector secreted by the Esx-3 secretion system as dimer with EsxG (Ilghari et al. 2011). Within the host macrophages, the dimer EsxH-EsxG targets Hrs (a component of the human endosomal sorting complex), impairing trafficking, and hindering phagosomal maturation and antigen presentation, thus contributing to the survival of the pathogen (Mehra et al. 2013, Portal-Celhay et al. 2016, Mittal et al. 2018). The observed signatures of selection could be due to the adaptation of L1 strains to human genotypes of *hrs* prevalent in South- and Southeast Asia. A similar signature of selection was observed in another Esx effector (EsxW), in MTBC strains belonging to L2 (Holt et al. 2018). Holt and colleagues found evidence for parallel evolution at one residue in the N-terminal loop of EsxW, outside the region covered by known epitopes, suggesting that the selective pressure was not due to antigen recognition.

The sampling effort in this study was considerable, and it provided a more complete picture compared to previous studies. Nevertheless, the sampling was not population-based, and for some regions the coverage was scarce (e.g. the Arabian Peninsula). Because of these limitations, our biogeographical analyses were limited to the subcontinental level. This approach revealed the global population structure and the main macroscopic patterns of diversity and migration of L1 and L3. However, MTBC populations are diverse also within subcontinental regions. For example, the MTBC population in Southern India is dominated by L1, while the most prevalent lineage in the North is L3 (Couvin et al. 2019). To investigate fine-scale processes in greater detail, including local adaptation, large population-based studies will be necessary.

In conclusion, the results presented here improve our knowledge about the TB epidemic around the Indian Ocean. A better understanding of the evolutionary dynamics of different MTBC populations might inform the development of control strategies for different regions. For example, *esxH* is part of several new vaccine candidates (Radošević et al. 2007, Abel et al. 2010; Hoang et al. 2013), and at least one of these, H4:IC31, is under clinical development in South Africa (Nemes et al. 2018; Bekker et al. 2020). In the light of our findings, and to develop a globally effective vaccine, it would be important to know if the results of the clinical trials in South Africa can be replicated in Southeast Asia, where there is a high prevalence of derived *esxH* haplotypes in the circulating MTBC populations.

## Methods

### Strain cultures, DNA extraction and genome sequencing

MTBC isolates previously identified as L1 and L3, either by SNP-typing or spoligotyping, were grown in Middlebrook 7H9 liquid medium supplemented with ADC and incubated at 37°C. Purified genomic DNA was obtained from cultures using a CTAB extraction method.

Whole genome sequencing was performed on libraries prepared from purified genomic DNA using Illumina Nextera ® XT library and NEBNext ® Ultra TM II FS DNA Library Prep Kits. Sequencing was performed using the Illumina HiSeq 2500 or NextSeq 500 paired-end technology.

The sequence data generated by this study has been deposited under the accession numbers PRJNA630228 and PRJXXXXX.

### Bioinformatic pipeline

We screened a large collection of publicly available whole genome sequences (Illumina) of MTBC strains belonging to L1 and L3, using the diagnostic SNPs described in Steiner et al. (2014). To cover geographic regions that were under-represented by this dataset, we additionally sequenced the genomes of 767 clinical strains (360 L1, and 407 L3).

For all samples, Illumina reads were trimmed with Trimmomatic v0.33 (SLIDINGWINDOW: 5:20,ILLUMINACLIP:{adapter}:2:30:10) (Bolger et al. 2014). Reads shorter than 20 bp were excluded from the downstream analysis. Overlapping paired-end reads were then merged with SeqPrep (overlap size = 15; https://github.com/jstjohn/SeqPrep). The resulting reads were mapped to the reconstructed MTBC ancestral sequence (Comas et al. 2013) with BWA v0.7.12 (mem algorithm; Li and Durbin 2010). Duplicated reads were marked by the MarkDuplicates module of Picard v 2.1.1 (https://github.com/broadinstitute/picard). The RealignerTargetCreator and IndelRealigner modules of GATK v.3.4.0 (McKenna et al. 2010) were used to perform local realignment of reads around Indels. Reads with alignment score lower than (0.93*read_length)-(read_length*4*0.07)) were excluded: this corresponds to more than 7 miss-matches per 100 bp.

SNPs were called with Samtools v1.2 mpileup (Li et al. 2009) and VarScan v2.4.1 (Koboldt et al. 2012) using the following thresholds: minimum mapping quality of 20, minimum base quality at a position of 20, minimum read depth at a position of 7X, minimum percentage of reads supporting the call 90%. SNPs in previously defined repetitive regions were excluded (i.e. PPE and PE-PGRS genes, phages, insertion sequences and repeats longer than 50 bp) as described before (Brites et al. 2019).

We applied the following filters: genomes were excluded if they had 1) an average coverage < 15x, 2) more than 50% of their SNPs excluded due to the strand bias filter, 3) more than 50% (or more than 1,000 in absolute number) of their SNPs having a percentage of reads supporting the call between 10% and 90%, 4) contained single nucleotide polymorphisms diagnostic for different MTBC lineages (Steiner et al. 2014), as this indicated that a mix of genomes was sequenced, 5) had more than 5,000 SNPs of difference compared to the reconstructed ancestral genome of the MTBC (Comas et al. 2013). Additionally, when multiple strains were sampled from the same patient, we retained only one. We further excluded all strains that had less SNPs than (average - (3 * standard deviation)) of the respective lineage (calculated after all previous filtering steps). We built SNP alignments for L1 and L3 separately, including only variable positions with less than 10% of missing data, and finally, we excluded all genomes with more than 10% of missing data in the alignment of the respective lineage. After all filtering steps, we were able to retrieve 4,968 strains with high quality genome sequences for further analyses (2,938 L1, 2,030 L3, Sup. Table 1).

### Analysis of sublineages

We used the curated datasets and inferred phylogenetic trees based on all polymorphic positions (excluding the ones in repetitive regions, see above) with raxml 8.2.11 (Stamatakis 2014; -m GTRCAT and -V options). We then identified sublineages following the classification (and using the diagnostic SNPs) of Coll et al. (2014).

### Molecular clock analyses with LSD

We selected all strains for which the year of sampling was known (2,499 strains, 1,672 L1, 827 L3). For both lineages, we built SNP alignments including only variable positions with less than 10% of missing data. We inferred phylogenetic trees with RAxML 8.2.11 (Stamatakis 2014), using the GTR model (-m GTRCAT -V options). Since the alignments contain only variable positions, we rescaled the branch lengths of the trees: rescaled_branch_length = ((branch_length * alignment_lengths) / (alignment_length + invariant_sites)). We rooted the trees using the genome sequence of a L2 strain as outgroup (accession number SAMEA4441446).

We used the least square method implemented in LSD v0.3-beta (To et al. 2015) to estimate the molecular clock rate with the QPD algorithm and calculating the confidence interval (options -f 100 and -s). We also performed a date randomization test by randomly reassigning the sampling dates among taxa 100 times and estimating the clock rate from the randomized and observed datasets. All LSD analyses were performed on the two lineages individually (L1 and L3), and on the five sublineages of L1.

### Molecular clock analyses with Beast

Bayesian molecular clock analyses are computationally demanding and they would be impossible to apply onto the complete datasets. Therefore, we sub-sampled the L1 and L3 datasets used for the LSD analysis to 400 genomes with two different strategies: 1) random subsampling 2) random subsampling keeping at least 25 genomes for each year of sampling (where possible; “weighted subsampling”). For this second subsampling strategy, we used Treemmer v0.3 (Menardo et al. 2018) with options -X 400, - pr, -lm, and -mc 25. For both subsampling schemes, we generated three subsets of the original datasets, resulting in six subsets for each lineage.

We assembled SNP alignments including only variable positions with less than 10% of missing data, and used jModelTest 2.1.10 v20160303 (Darriba et al. 2012) to identify the best fitting nucleotide substitution model (according to the Akaike information criterion) among 11 possible schemes including unequal nucleotide frequencies (total models = 22, options -s 11 and -f).

We performed Bayesian inference with Beast 2.5 (Bouckaert et al. 2019). We corrected the xml files to specify the number of invariant sites as indicated here: https://groups.google.com/forum/#!topic/beast-users/QfBHMOqImFE, and used the tip sampling year as calibration. We assumed the best fitting nucleotide substitution model as identified by jModelTest, a relaxed lognormal clock model (Drummond et al. 2006) and an exponential population size coalescent prior. We chose a 1/x prior for the population size [0–10^9^], a 1/x prior for the mean of the lognormal distribution of the clock rate [10−^10^–10−^5^], a normal(0,1) prior for the standard deviation of the lognormal distribution of the clock rate [0 –infinity]. For the exponential growth rate prior, we used the standard Laplace distribution [-infinity–infinity]. For all datasets, we ran at least two runs, we used Tracer 1.7.1 (Rambaut et al. 2018) to identify and exclude the burn-in, to evaluate convergence among runs and to calculate the estimated sample size. We stopped the runs when at least two chains reached convergence, and the ESS of the posterior and of all parameters were larger than 200.

Since we detected a strong temporal signal only for L3, we performed a set of additional analyses of the subsets of L3. We repeated the Beast analysis with an extended Bayesian Skyline Plot (BSP) prior instead of the exponential growth prior, and performed a nested sampling analysis (Russel et al. 2019) to identify which of these two models (exponential growth and extended BSP) fitted the data best. The nested sampling was run with chainLength = 20000, particleCount= 4, and subChainLength = 10000.

All xml input files are available as Supplementary Files.

### Datasets for biogeography analyses

For the biogeography analysis, we considered only genome sequences obtained from strains for which the locality of sampling was known. When the country of sampling did not correspond to the country of origin of the patient (or was unknown), we considered as sampling locality the country of origin of the patient (this affected 187 strains, 121 L1 and 66 L3). Furthermore, similarly to other studies (O’Neill et al. 2019), we excluded all genomes that were sampled from Europe, North America and Australia, as most contemporary infections in these regions affect recent migrants. The final dataset for the biogeography analyses comprised 3,082 strains (2,061 L1 and 1,021 L3). We assigned the different isolates to different subcontinental geographic regions according to sampling locality. To do this, we followed a modified version of the United Nations geographic scheme (https://unstats.un.org/unsd/methodology/m49/; Sup. Table 1 and Figs. 1-3).

### Phylogeography analysis with PASTML

For both lineages, we built SNP alignments including only variable positions with less than 10% of missing data, and all strains with known sampling locality (excluding strains from North America, Europe and Australia; see above, 2,061 L1 genomes and 1,021 L3 genomes). We inferred phylogenetic trees with raxml 8.2.11 (Stamatakis 2014), using the GTR model (-m GTRCAT -V options). Since the alignments contain only variable positions, we re-scaled the branch lengths of the trees: rescaled_branch_length = ((branch_length * alignment_lengths) / (alignment_length + invariant_sites)).

Since PASTML needs a time tree as input, we calibrated the phylogenies with LSD, assuming a clock rate of 1.4×10^−7^ for L1, and 9×10^−8^ for L3. In this analysis, genomes for which the sampling date was not known were assumed to have been sampled between 1995 and 2018, which is the period in which all strains with known date of isolation were sampled. Importantly, using different clock rates for this analysis would only change the time scale of the trees, but not the reconstruction of the ancestral characters.

We assigned to each strain the subcontinental geographic region of origin as character, and used PASTML (Ishikawa et al. 2019) to reconstruct the ancestral geographical ranges and their changes along the trees of L1 and L3. We used the MPPA as prediction method (standard settings) and added the character predicted by the joint reconstruction even if it was not selected by the Brier score (option - forced_joint). Additionally, we repeated the PASTML analysis for the sublineages of L1 individually.

### Phylogeography analysis with Mascot

As a complementary method to reconstruct the ancestral range and the migration pattern of different populations, we used the Beast package Mascot (Müller et al. 2018). We assumed that strains sampled in the different subcontinental regions represent distinct subpopulations, and we considered only populations for which we had at least 75 genome sequences: four populations for L1 (East Africa, South Asia, Southeast Asia (mainland) and Southeast Asia (islands)), and two populations for L3 (East Africa and South Asia). For computational reasons, we subsampled the two datasets (L1 and L3) to ~300 strains. We sampled an equal number of strains from each geographic region (where possible), and within regions, we randomly sampled an equal number of strains from each country (where possible). This sub-sampling scheme resulted in two subsets of 303 and 300 strains for L1 and L3, respectively.

We assembled SNP alignments including only variable positions with less than 10% of missing data, and used jModelTest 2.1.10 v20160303 (Darriba et al. 2012) to identify the best fitting nucleotide substitution model as described above.

We performed Bayesian inference with Beast2.5 (Bouckaert et al. 2019). We corrected the xml files to specify the number of invariant sites as indicated here: https://groups.google.com/forum/#!topic/beast-users/QfBHMOqImFE. We assumed a lognormal uncorrelated clock and we fixed the mean of the lognormal distribution of the clock rate to 1.4×10^−7^ (L1) and 9×10^−8^ (L3). We assigned the tip sampling years to the different strains, and when the sampling time was unknown, we assumed a uniform prior from 1995 to 2018 (similarly to what done in the PASTML analysis). We further assumed the best fitting nucleotide substitution model as identified by jModelTest, a gamma prior for the standard deviation of the lognormal distribution of the clock rate [0 –infinity], and a lognormal prior for the population size with standard deviation = 0.2, and mean estimated in real space. Finally, we used an exponential distribution with mean = 10^−4^ as prior for the migration rates. For each analysis, we ran at least two runs. We used Tracer 1.7.1 (Rambaut et al. 2018) to identify and exclude the burn-in, to evaluate convergence among runs, and to calculate the estimated sample size. We stopped the runs when at least two chains reached convergence, and the ESS of the posterior, prior and of the parameters of interest (population sizes and migration rates) were larger than 200.

The xml input files are available as Supplementary Files.

### L2 and L4 datasets

For the analysis of *esxH*, we wanted to compare the results obtained for L1 and L3 with the other two major human-adapted MTBC lineages L2 and L4. Therefore, we compiled two datasets of publicly available genome sequences for these two lineages. We applied the same bioinformatic pipeline described above: genomes were excluded if they had 1) an average coverage < 15x, 2) more than 50% (or more than 1,000 in absolute number) of their SNPs having a percentage of reads supporting the call between 10% and 90%, or 3) contained single nucleotide polymorphisms diagnostic for different MTBC lineages (Steiner et al. 2014), as this could indicate mixed infection. The final dataset consisted of 6,752 L2 genome sequences (with 140,309 polymorphic positions) and 10,466 L4 genome sequences (with 277,648 polymorphic positions; Sup. Table 1). We reconstructed the phylogenetic tree of L2 with raxml 8.2.11 (Stamatakis 2014) as described above. Due to the large size of the dataset, we used FastTree (Price et al. 2010) with options -nt -nocat -nosupport to reconstruct the phylogenetic tree of L4.

### Epitopes analysis

We downloaded the amino acid sequence of all MTBC epitopes described for *Homo sapiens* from the immune epitope database (https://www.iedb.org/; downloaded on the 03.08.2020). We considered separately MHCI epitopes and MHCII epitopes. We mapped the epitope sequences onto the H37Rv genome (GCF_000195955.2) using tblastn and excluded sequences mapping equally well to multiple loci in the H37Rv genome. Additionally, we retained only epitopes that mapped with two mismatches or less over the whole epitope length. This resulted in a final list of 539 MHCI epitopes, and 1,144 MHCII epitopes. (Sup. Table 7). We used the datasets obtained for the biogeography analysis (2,061 genome sequences for L1, and 1,021 genome sequences for L3).

For each lineage, we independently assembled a multiple sequence alignment for each epitope. We then translated the sequence to amino acids and used PAUP 4.a (Wilgenbusch and Swofford. 2003) to reconstruct the replacement history of all polymorphic positions on the rooted phylogenetic trees. We used two maximum parsimony algorithms (ACCTRAN and DELTRAN) and considered only the events reconstructed by both algorithms.

### Analysis of *esxH*

We considered the first 18 amino acids of *esxH* (MSQIMYNYPAMLGHAGDM), which was by far the epitope with most amino acid replacements in L1. We expanded the analysis of this epitope to the L2 and L4 datasets, so that we could compare the results with L1 and L3. For the PAML analysis, we randomly selected 500 strains from each MTBC lineage. We used the phylogenetic tree reconstructed by RAxML (same settings as above), and the gene alignment to estimate the branch lengths of the tree using the M0 codon model implemented in PAML 4.9e (Yang 2007). This step was necessary to obtain a tree with the branch length in expected substitutions per codon. We then fitted two alternative codon models (M1a and M2a) to the trees and alignments. M1a allows ω to be variable across sites, and it assumes two different ω (0 < ω_0_ <1, and ω_1_ = 1), modeling nearly neutral evolution; M2a assumes one additional ω (ω_2_ > 1) compared to M1a, thus modeling positive selection. We performed a likelihood ratio test between the two models as described in Zhang et al. (2005). Templates for the control files of the codeml analyses (M0, M1a, M2a) are available as Supplementary Files. The codon under positive selection were identified with the Bayes empirical Bayes method (Yang et al. 2005).

To test whether the derived haplotypes of this epitope were associated with a specific geographic region, we constructed a statistical test analogous to PhyC (Farhat et al. 2013), which is normally used to test for association between a variant and phenotypic drug resistance. We simulated mutations on the phylogenetic tree of L1 and counted how many strains from each region resulted to have a derived state according to the simulation. Under this procedure, mutations occur randomly on the tree, and therefore independently from the geographic region where the tips were sampled. For each test, we performed 10,000 simulations to obtain a null distribution, and we compared this distribution to the observed data to obtain empirical p-values. We used the same geographic region used for the biogeography analysis. Additionally, we considered East Africa excluding Madagascar.

R code and input files to perform this test are available as Supplementary Files.

### Prediction of binding affinity between T cell epitopes and HLA alleles

We considered three HLA loci: HLA-A, HLA-B and HLA-DRB1. We used the allele frequency database (Gonzalez-Galarza et al. 2020) to identify alleles that are prevalent in East Africa and not in South Asia and Southeast Asia, or the other way around. Because the coverage of the allele frequency database is patchy, we focused on the following countries: Kenya and Zimbabwe as representatives of East Africa; India, Thailand and Taiwan as representatives of South- and Southeast Asia. We identified:

1. Alleles that had a frequency of 10% or more in at least one population in South- and Southeast Asia, but had frequencies lower than 10% in all East African populations.
2. Alleles that had a frequency of 10% or more in at least one population in East Africa, but had frequencies lower than 10% in all populations in South- and Southeast Asia.
3. Alleles that had a frequency of 10% or more in at least one population both in South- and Southeast Asia and in East Africa. For all these alleles, we performed *in silico* binding prediction with three epitopes: the ancestral epitope at the *esxH* N terminus (MSQIMYNYPAMLGHAGDM), and the two most frequently observed derived epitopes (MSQIMYNYPTMLGHAGDM, and MSQIMYNYPVMLGHAGDM). For HLA-A and HLA-B alleles, we used the NetMHCPan4.1 server (Reynisson et al. 2020) with standard settings. For HLA-DRB1 alleles, we used the prediction tool of the immune epitope database (https://www.iedb.org/).

### Genome wide scan for positive selection with PAML

For this analysis, we used the subsets generated for the Mascot analysis. These datasets are representative of the populations of L1 and L3 in their core geographic ranges, and are computationally treatable. We generated sequence alignments for all genes individually, excluding genes in repetitive regions of the genome (see above). Because some genes are deleted in L1 but not in L3, or the other way around, we obtained a slightly different number of gene alignments for the two lineages (L1: 3,623, L3: 3,622). For each gene, we performed a test for positive selection with PAML as described above for *esxH*. We considered as under positive selection all genes, for which the likelihood ratio test resulted in a Bonferroni-corrected p-value < 0.05.

### Drug resistance mutations profiles

We considered 196 mutations conferring resistance to different antibiotics (Payne et al. 2019). We extracted the respective genomic positions from the vcf file of the 4,968 genomes of the complete curated data set (2,938 L1, 2,030 L3) and assembled them in phylip format. To determine the number of independent mutations, we reconstructed the nucleotide changes on the phylogenetic tree rooted with the L2 strain (SAMEA4441446). To do this, we used the maximum parsimony ACCTRAN and DELTRAN algorithms implemented in PAUP 4.0a (Wilgenbusch and Swofford. 2003), and considered only the events reconstructed by both algorithms.

## Acknowledgements

This work was supported by the Swiss National Science Foundation (grants 310030_188888, CRSII5_177163, IZRJZ3_164171 and IZLSZ3_170834) and the European Research Council (309540‑EVODRTB and 883582-ECOEVODRTB). Calculations were performed at sciCORE (http://scicore.unibas.ch/) scientific computing core facility at the University of Basel.

## Supplementary Files

Supplementary results and code are available here: https://github.com/fmenardo/MTBC_L1_L3.

## References

Abel, B., Tameris, M., Mansoor, N., Gelderbloem, S., Hughes, J., Abrahams, D., … & Hawkridge, A. (2010). The novel tuberculosis vaccine, AERAS-402, induces robust and polyfunctional CD4+ and CD8+ T cells in adults. American journal of respiratory and critical care medicine, 181(12), 1407–1417.

Allen, R. B. (2013). Indian Ocean transoceanic migration, 16th–19th century. The Encyclopedia of Global Human Migration.

Bekker, L. G., Dintwe, O., Fiore-Gartland, A., Middelkoop, K., Hutter, J., Williams, A., … & DiazGranados, C. A. (2020). A phase 1b randomized study of the safety and immunological responses to vaccination with H4: IC31, H56: IC31, and BCG revaccination in Mycobacterium tuberculosis-uninfected adolescents in Cape Town, South Africa. EClinicalMedicine, 100313.

Blench, R. (2010). Evidence for the Austronesian voyages in the Indian Ocean. The global origins and development of seafaring, 239, 48.

Boivin, N., Crowther, A., Helm, R., & Fuller, D. Q. (2013). East Africa and Madagascar in the Indian Ocean world. Journal of World Prehistory, 26(3), 213–281.

Bolger, A. M., Lohse, M., & Usadel, B. (2014). Trimmomatic: a flexible trimmer for Illumina sequence data. Bioinformatics, 30(15), 2114–2120.

Bos, K. I., Harkins, K. M., Herbig, A., Coscolla, M., Weber, N., Comas, I., … & Campbell, T. J. (2014). Pre-Columbian mycobacterial genomes reveal seals as a source of New World human tuberculosis. Nature, 514(7523), 494–497.

Bottai, D., Frigui, W., Sayes, F., Di Luca, M., Spadoni, D., Pawlik, A., … & Mangenot, S. (2020). TbD1 deletion as a driver of the evolutionary success of modern epidemic Mycobacterium tuberculosis lineages. Nature communications, 11(1), 1–14.

Bouckaert, R., Vaughan, T. G., Barido-Sottani, J., Duchêne, S., Fourment, M., Gavryushkina, A., … & Drummond, A. J. (2019). BEAST 2.5: An advanced software platform for Bayesian evolutionary analysis. PLoS computational biology, 15(4), e1006650.

Brahmajothi, V., Pitchappan, R. M., Kakkanaiah, V. N., Sashidhar, M., Rajaram, K., Ramu, S., … & Prabhakar, R. (1991). Association of pulmonary tuberculosis and HLA in south India. Tubercle, 72(2), 123–132.

Brites, D., Loiseau, C., Menardo, F., Borrell, S., Boniotti, M. B., Warren, R., … & Fyfe, J. A. (2018). A new phylogenetic framework for the animal-adapted Mycobacterium tuberculosis complex. Frontiers in microbiology, 9, 2820.

Brucato, N., Kusuma, P., Cox, M. P., Pierron, D., Purnomo, G. A., Adelaar, A., … & Ricaut, F. X. (2016). Malagasy genetic ancestry comes from an historical Malay trading post in Southeast Borneo. Molecular biology and evolution, 33(9), 2396–2400.

Brucato, N., Fernandes, V., Mazières, S., Kusuma, P., Cox, M. P., Wainaina Ng’ang’a, J., … & Fin, B. (2018). The Comoros show the earliest Austronesian gene flow into the Swahili corridor. The American Journal of Human Genetics, 102(1), 58–68.

Brucato, N., Fernandes, V., Kusuma, P., Černý, V., Mulligan, C. J., Soares, P., … & Cox, M. P. (2019). Evidence of Austronesian genetic lineages in East Africa and South Arabia: complex dispersal from Madagascar and Southeast Asia. Genome Biology and Evolution, 11(3), 748–758.

Chihota, V. N., Niehaus, A., Streicher, E. M., Wang, X., Sampson, S. L., Mason, P., … & Kasongo, W. (2018). Geospatial distribution of Mycobacterium tuberculosis genotypes in Africa. PLoS One, 13(8), e0200632.

Coll, F., McNerney, R., Guerra-Assuncao, J. A., Glynn, J. R., Perdigao, J., Viveiros, M., … & Clark, T. G. (2014). A robust SNP barcode for typing Mycobacterium tuberculosis complex strains. Nature communications, 5(1), 1–5.

Comas, I., Chakravartti, J., Small, P. M., Galagan, J., Niemann, S., Kremer, K., … & Gagneux, S. (2010). Human T cell epitopes of Mycobacterium tuberculosis are evolutionarily hyperconserved. Nature genetics, 42(6), 498–503.

Comas, I., Coscolla, M., Luo, T., Borrell, S., Holt, K. E., Kato-Maeda, M., … & Yeboah-Manu, D. (2013). Out-of-Africa migration and Neolithic coexpansion of Mycobacterium tuberculosis with modern humans. Nature genetics, 45(10), 1176–1182.

Conceição, E. C., Refregier, G., Gomes, H. M., Olessa-Daragon, X., Coll, F., Ratovonirina, N. H., … & Gagneux, S. (2019). Mycobacterium tuberculosis lineage 1 genetic diversity in Pará, Brazil, suggests common ancestry with east-African isolates potentially linked to historical slave trade. Infection, Genetics and Evolution, 73, 337–341.

Coscolla, M., Copin, R., Sutherland, J., Gehre, F., de Jong, B., Owolabi, O., … & Gagneux, S. (2015). M. tuberculosis T cell epitope analysis reveals paucity of antigenic variation and identifies rare variable TB antigens. Cell host & microbe, 18(5), 538–548.

Couvin, D., Reynaud, Y., & Rastogi, N. (2019). Two tales: Worldwide distribution of Central Asian (CAS) versus ancestral East-African Indian (EAI) lineages of Mycobacterium tuberculosis underlines a remarkable cleavage for phylogeographical, epidemiological and demographical characteristics. PloS one, 14(7), e0219706.

Darriba, D., Taboada, G. L., Doallo, R., & Posada, D. (2012). jModelTest 2: more models, new heuristics and parallel computing. Nature methods, 9(8), 772–772.

De Jong, B. C., Antonio, M., & Gagneux, S. (2010). Mycobacterium africanum—review of an important cause of human tuberculosis in West Africa. PLoS Negl Trop Dis, 4(9), e744.

Devis, M. C., Brites, D., Menardo, F., Loiseau, C. M., Otchere, I. D., Asante-Poku, A., … & Dissou, A. (2020). Phylogenomics of Mycobacterium africanum reveals a new lineage and a complex evolutionary history. bioRxiv.

Dou, H. Y., Chen, Y. Y., Kou, S. C., & Su, I. J. (2015). Prevalence of Mycobacterium tuberculosis strain genotypes in Taiwan reveals a close link to ethnic and population migration. Journal of the Formosan Medical Association, 114(6), 484–488.

Drummond, A. J., Ho, S. Y., Phillips, M. J., & Rambaut, A. (2006). Relaxed phylogenetics and dating with confidence. PLoS Biol, 4(5), e88.

Duchêne, S., Holt, K. E., Weill, F. X., Le Hello, S., Hawkey, J., Edwards, D. J., … & Holmes, E. C. (2016). Genome-scale rates of evolutionary change in bacteria. Microbial genomics, 2(11).

Farhat, M. R., Shapiro, B. J., Kieser, K. J., Sultana, R., Jacobson, K. R., Victor, T. C., … & Kaur, D. (2013). Genomic analysis identifies targets of convergent positive selection in drug-resistant Mycobacterium tuberculosis. Nature genetics, 45(10), 1183–1189.

Fenner, L., Egger, M., Bodmer, T., Altpeter, E., Zwahlen, M., Jaton, K., … & Gagneux, S. (2012). Effect of mutation and genetic background on drug resistance in Mycobacterium tuberculosis. Antimicrobial agents and chemotherapy, 56(6), 3047–3053

Gagneux, S. (Ed.). (2017). Strain variation in the Mycobacterium tuberculosis complex: its role in biology, epidemiology and control (Vol. 1019). Springer.

Gagneux, S. (2018). Ecology and evolution of Mycobacterium tuberculosis. Nature Reviews Microbiology, 16(4), 202.

Glaziou, P. (2020). Predicted impact of the COVID-19 pandemic on global tuberculosis deaths in 2020. medRxiv.

Gonzalez-Galarza, F. F., McCabe, A., Santos, E. J. M. D., Jones, J., Takeshita, L., Ortega-Rivera, N. D., … & Middleton, D. (2020). Allele frequency net database (AFND) 2020 update: gold-standard data classification, open access genotype data and new query tools. Nucleic Acids Research, 48(D1), D783–D788.

Ho, S. Y., Phillips, M. J., Cooper, A., & Drummond, A. J. (2005). Time dependency of molecular rate estimates and systematic overestimation of recent divergence times. Molecular biology and evolution, 22(7), 1561–1568.

Hoang, T., Aagaard, C., Dietrich, J., Cassidy, J. P., Dolganov, G., Schoolnik, G. K., … & Andersen, P. (2013). ESAT-6 (EsxA) and TB10. 4 (EsxH) based vaccines for pre-and post-exposure tuberculosis vaccination. PloS one, 8(12), e80579.

Hogan, A. B., Jewell, B. L., Sherrard-Smith, E., Vesga, J. F., Watson, O. J., Whittaker, C., … & Baguelin, M. (2020). Potential impact of the COVID-19 pandemic on HIV, tuberculosis, and malaria in low-income and middle-income countries: a modelling study. The Lancet Global Health, 8(9), e1132–e1141.

Holt, K. E., McAdam, P., Thai, P. V. K., Thuong, N. T. T., Ha, D. T. M., Lan, N. N., … & Thwaites, G. (2018). Frequent transmission of the Mycobacterium tuberculosis Beijing lineage and positive selection for the EsxW Beijing variant in Vietnam. Nature genetics, 50(6), 849–856.

Ilghari, D., Lightbody, K. L., Veverka, V., Waters, L. C., Muskett, F. W., Renshaw, P. S., & Carr, M. D. (2011). Solution Structure of the Mycobacterium tuberculosis EsxG· EsxH Complex functional implications and comparisons with other m. tuberculosis esx family complexes. Journal of Biological Chemistry, 286(34), 29993–30002.

Ishikawa, S. A., Zhukova, A., Iwasaki, W., & Gascuel, O. (2019). A fast likelihood method to reconstruct and visualize ancestral scenarios. Molecular biology and evolution, 36(9), 2069–2085.

Koboldt, D. C., Zhang, Q., Larson, D. E., Shen, D., McLellan, M. D., Lin, L., … & Wilson, R. K. (2012). VarScan 2: somatic mutation and copy number alteration discovery in cancer by exome sequencing. Genome research, 22(3), 568–576.

Li, H., Handsaker, B., Wysoker, A., Fennell, T., Ruan, J., Homer, N., … & Durbin, R. (2009). The sequence alignment/map format and SAMtools. Bioinformatics, 25(16), 2078–2079.

Li, H., & Durbin, R. (2010). Fast and accurate long-read alignment with Burrows–Wheeler transform. Bioinformatics, 26(5), 589–595.

McKenna, A., Hanna, M., Banks, E., Sivachenko, A., Cibulskis, K., Kernytsky, A., … & DePristo, M. A. (2010). The Genome Analysis Toolkit: a MapReduce framework for analyzing next-generation DNA sequencing data. Genome research, 20(9), 1297–1303.

McQuaid, C. F., McCreesh, N., Read, J. M., Sumner, T., Houben, R. M., White, R. G., … & CMMID COVID-19 Working Group. (2020). The potential impact of COVID-19-related disruption on tuberculosis burden. European Respiratory Journal, 56(2).

Mehra, A., Zahra, A., Thompson, V., Sirisaengtaksin, N., Wells, A., Porto, M., … & Rogan, D. (2013). Mycobacterium tuberculosis type VII secreted effector EsxH targets host ESCRT to impair trafficking. PLoS Pathog, 9(10), e1003734.

Menardo, F., Loiseau, C., Brites, D., Coscolla, M., Gygli, S. M., Rutaihwa, L. K., … & Gagneux, S. (2018). Treemmer: a tool to reduce large phylogenetic datasets with minimal loss of diversity. BMC bioinformatics, 19(1), 1–8.

Menardo, F., Duchêne, S., Brites, D., & Gagneux, S. (2019). The molecular clock of Mycobacterium tuberculosis. PLoS pathogens, 15(9), e1008067.

Mittal, E., Skowyra, M. L., Uwase, G., Tinaztepe, E., Mehra, A., Köster, S., … & Philips, J. A. (2018). Mycobacterium tuberculosis type VII secretion system effectors differentially impact the ESCRT endomembrane damage response. MBio, 9(6).

Müller, N. F., Rasmussen, D., & Stadler, T. (2018). MASCOT: parameter and state inference under the marginal structured coalescent approximation. Bioinformatics, 34(22), 3843–3848.

Nemes, E., Geldenhuys, H., Rozot, V., Rutkowski, K. T., Ratangee, F., Bilek, N., … & Mulenga, H. (2018). Prevention of M. tuberculosis infection with H4: IC31 vaccine or BCG revaccination. New England Journal of Medicine, 379(2), 138–149.

Ngabonziza, J. C. S., Loiseau, C., Marceau, M., Jouet, A., Menardo, F., Tzfadia, O., … & Diels, M. (2020). A sister lineage of the Mycobacterium tuberculosis complex discovered in the African Great Lakes region. Nature Communications, 11(1), 1–11.

O’Neill, M. B., Shockey, A., Zarley, A., Aylward, W., Eldholm, V., Kitchen, A., & Pepperell, C. S. (2019). Lineage specific histories of Mycobacterium tuberculosis dispersal in Africa and Eurasia. Molecular ecology, 28(13), 3241–3256.

Payne, J. L., Menardo, F., Trauner, A., Borrell, S., Gygli, S. M., Loiseau, C., … & Hall, A. R. (2019). Transition bias influences the evolution of antibiotic resistance in Mycobacterium tuberculosis. PLoS biology, 17(5), e3000265.

Pepperell, C. S., Casto, A. M., Kitchen, A., Granka, J. M., Cornejo, O. E., Holmes, E. C., … & Feldman, M. W. (2013). The role of selection in shaping diversity of natural M. tuberculosis populations. PLoS Pathog, 9(8), e1003543.

Peters, J. S., Ismail, N., Dippenaar, A., Ma, S., Sherman, D. R., Warren, R. M., & Kana, B. D. (2020). Genetic Diversity in Mycobacterium tuberculosis Clinical Isolates and Resulting Outcomes of Tuberculosis Infection and Disease. Annual Review of Genetics, 54.

Pierron, D., Razafindrazaka, H., Pagani, L., Ricaut, F. X., Antao, T., Capredon, M., … & Letellier, T. (2014). Genome-wide evidence of Austronesian–Bantu admixture and cultural reversion in a hunter-gatherer group of Madagascar. Proceedings of the National Academy of Sciences, 111(3), 936–941.

Portal-Celhay, C., Tufariello, J. M., Srivastava, S., Zahra, A., Klevorn, T., Grace, P. S., … & Philips, J. A. (2016). Mycobacterium tuberculosis EsxH inhibits ESCRT-dependent CD4+ T-cell activation. Nature microbiology, 2(2), 1–9.

Price, M. N., Dehal, P. S., & Arkin, A. P. (2010). FastTree 2–approximately maximum-likelihood trees for large alignments. PloS one, 5(3), e9490.

Qu, H. Q., Fisher-Hoch, S. P., & McCormick, J. B. (2011). Knowledge gaining by human genetic studies on tuberculosis susceptibility. Journal of human genetics, 56(3), 177–182.

Rabahi, M. F., Conceição, E. C., de Paiva, L. O., Souto, M. V. M. L., Sisco, M. C., de Waard, J., … & Campos, C. E. D. (2020). Characterization of Mycobacterium tuberculosis var. africanum isolated from a patient with pulmonary tuberculosis in Brazil. Infection, Genetics and Evolution, 85, 104550.

Radošević, K., Wieland, C. W., Rodriguez, A., Weverling, G. J., Mintardjo, R., Gillissen, G., … & van der Poll, T. (2007). Protective immune responses to a recombinant adenovirus type 35 tuberculosis vaccine in two mouse strains: CD4 and CD8 T-cell epitope mapping and role of gamma interferon. Infection and immunity, 75(8), 4105–4115.

Rambaut, A., Drummond, A. J., Xie, D., Baele, G., & Suchard, M. A. (2018). Posterior summarization in Bayesian phylogenetics using Tracer 1.7. Systematic biology, 67(5), 901.

Reynisson, B., Alvarez, B., Paul, S., Peters, B., & Nielsen, M. (2020). NetMHCpan-4.1 and NetMHCIIpan-4.0: improved predictions of MHC antigen presentation by concurrent motif deconvolution and integration of MS MHC eluted ligand data. Nucleic Acids Research.

Russel, P. M., Brewer, B. J., Klaere, S., & Bouckaert, R. R. (2019). Model selection and parameter inference in phylogenetics using Nested Sampling. Systematic biology, 68(2), 219–233.

Salie, M., van der Merwe, L., Möller, M., Daya, M., van der Spuy, G. D., van Helden, P. D., … & Hoal, E. G. (2014). Associations between human leukocyte antigen class I variants and the Mycobacterium tuberculosis subtypes causing disease. The Journal of infectious diseases, 209(2), 216–223.

Saunders, M. J., & Evans, C. A. (2020). COVID-19, tuberculosis, and poverty: preventing a perfect storm. European Respiratory Journal; DOI: 10.1183/13993003.01348-2020

Smittipat, N., Miyahara, R., Juthayothin, T., Billamas, P., Dokladda, K., Imsanguan, W., … & Wongjai, J. (2019). Indo-Oceanic Mycobacterium tuberculosis strains from Thailand associated with higher mortality. The International Journal of Tuberculosis and Lung Disease, 23(9), 972–979.

Stamatakis, A. (2014). RAxML version 8: a tool for phylogenetic analysis and post-analysis of large phylogenies. Bioinformatics, 30(9), 1312–1313.

Steiner, A., Stucki, D., Coscolla, M., Borrell, S., & Gagneux, S. (2014). KvarQ: targeted and direct variant calling from fastq reads of bacterial genomes. BMC genomics, 15(1), 881.

Sveinbjornsson, G., Gudbjartsson, D. F., Halldorsson, B. V., Kristinsson, K. G., Gottfredsson, M., Barrett, J. C., … & Helgadottir, H. T. (2016). HLA class II sequence variants influence tuberculosis risk in populations of European ancestry. Nature genetics, 48(3), 318–322.

To, T. H., Jung, M., Lycett, S., & Gascuel, O. (2015). Fast dating using least-squares criteria and algorithms. Systematic biology, 65(1), 82–9

Vejbaesya, S., Chierakul, N., Luangtrakool, K., Srinak, D., & Stephens, H. A. F. (2002). Associations of HLA class II alleles with pulmonary tuberculosis in Thais. European journal of immunogenetics, 29(5), 431–434.

White, Z., Painter, J., Douglas, P., Abubakar, I., Njoo, H., Archibald, C., … & Posey, D. L. (2017). Immigrant arrival and tuberculosis among large immigrant-and refugee-receiving countries, 2005–2009. Tuberculosis Research and Treatment, 2017.

Wiens, K. E., Woyczynski, L. P., Ledesma, J. R., Ross, J. M., Zenteno-Cuevas, R., Goodridge, A., … & Ray, S. E. (2018). Global variation in bacterial strains that cause tuberculosis disease: a systematic review and meta-analysis. BMC medicine, 16(1), 1–13.

Wilgenbusch, J. C., & Swofford, D. (2003). Inferring evolutionary trees with PAUP. Current protocols in bioinformatics, (1), 6–4.

World Health Organization. (2020). Global Tuberculosis Report (https://apps.who.int/iris/bitstream/handle/10665/336069/9789240013131-eng.pdf?ua=1)

Yang, Z., Wong, W. S., & Nielsen, R. (2005). Bayes empirical Bayes inference of amino acid sites under positive selection. Molecular biology and evolution, 22(4), 1107–1118.

Yang, Z. (2007). PAML 4: phylogenetic analysis by maximum likelihood. Molecular biology and evolution, 24(8), 1586–1591.

Yuliwulandari, R., Sachrowardi, Q., Nakajima, H., Kashiwase, K., Hirayasu, K., Mabuchi, A., … & Tokunaga, K. (2010). Association of HLA-A,-B, and-DRB1 with pulmonary tuberculosis in western Javanese Indonesia. Human immunology, 71(7), 697–701.

Zhang, J., Nielsen, R., & Yang, Z. (2005). Evaluation of an improved branch-site likelihood method for detecting positive selection at the molecular level. Molecular biology and evolution, 22(12), 2472–2479.

