## Supplementary Information for "Local adaptation in populations of *Mycobacterium tuberculosis* endemic to the Indian Ocean Rim"

**Phylogeography of L1 and L3**

The goal of this analysis was to reconstruct the historical spread of L1 and L3 through the world, and the patterns of migration among different regions. We considered the genome sequence of 2,061 L1 strains originating from 42 countries in 10 geographic regions, and 1,021 L3 strains originating from 32 countries in 8 geographic regions (Figs. 1-2). We assigned the subcontinental geographic region of origin to each strain, and used PASTML (Ishikawa et al. 2019) to reconstruct the ancestral geographical ranges, and their changes along the trees.

For both lineages PASTML inferred South Asia as the most likely range of the most recent common ancestor (MRCA). Moreover, we found a striking pattern of migration out of South Asia toward all other regions. We detected several recurring migrations from South Asia to East Africa, followed by further dispersion to the other African regions (South Africa, West and Central Africa, North Africa). We also identified additional migration events between different regions, but almost no migration back to South Asia (Figs. 2-3; Sup Files 1-2).

We repeated the PASTML analysis on the individual sublineages of L1, the results confirmed the prominent role of South Asia in the global dispersion of L1. Additionally, in contrast to what obtained with the complete dataset, this analysis inferred a more important role of Southeast Asia in the spread of L1, with several migration events between the islands and mainland of Southeast Asia, and from Southeast Asia towards other regions (Sup. Fig. 7).

To evaluate the robustness of these results we adopted a different approach based on coalescent theory, which allow to estimate migration rates between different populations and population sizes. To do this we used Mascot (Müller et al. 2018). Compared to PASTML, Mascot is not only based on a different modeling framework, but it implements a Bayesian approach that account for the uncertainty in the parameter estimations, including the tree topology and branch lengths. However, this comes with some limitations: Mascot can estimate the population sizes and migration rates reliably, only if there are enough coalescent events within and between population respectively. Therefore, we assumed that each geographic region (the same used for the PASTML analysis) represented a single population, and considered only populations for which we had the genome sequence of at least 75 strains (4 populations for L1: East Africa, South Asia, Southeast Asia (mainland), and Southeast Asia (islands); 2 populations for L3: East Africa and South Asia). For computational reasons we sub-sampled the two datasets (L1 and L3) to ~300 strains (see Methods for details).

Mascot inferred South Asia as the range of the MRCA of both L1 and L3, additionally we found strongly asymmetric migration rates: the (forward) migration rates from South Asia to all other regions were 3 to 26 times larger than the rates of migration toward South Asia or between the other regions (Sup. Table 2, Figs. 2-3). Finally, the effective population sizes of the four L1 populations were similar, while for L3, the South Asian population was estimated to have an effective size more than twice the effective size of the East African population (Sup. Table 2, Figs. 2-3).

In summary, we found that South Asia was the most likely geographic range of the MRCAs of L1 and L3, and that the spread of the two lineages was characterized by asymmetric migration from South Asia to the rest of the world, with little migration back to South Asia. Both methods (PASTML and Mascot) have limitations, however they are based on different models, and they use different statistical machinery. The fact that such different methods produced the same results support their robustness to different assumptions.

**Molecular clock analyses of L1**

For the molecular clock analyses, we considered all genome sequences for which the year of isolation was known (1,672 genome sequences). We used the inferred phylogenetic tree and the years of isolation of the tips to time-calibrate the tree with LSD (To et al. 2015), and we performed a date randomization test (DRT) with 100 randomized replicates (see Methods for details). We found a very weak temporal signal: the estimate obtained with the observed data was barely distinguishable from the ones obtained with the randomized replicates (Sup. Table 3, Sup. Fig 8). Additionally, the lower limit of the 95% CI reached the minimum value allowed by LSD, further indicating a lack of temporal structure. We repeated this analysis on the five sublineages of L1 individually, and found that three of them did not pass the DRT , showing a lack of temporal signal (L1.1.1, L1.1.3, and L1.2.1), while the remaining two (L1.1.2 and L1.2.2) had a weak to moderate temporal signal (Sup. Fig. 9). For the two sublineages with some temporal signal, the clock rate estimates were ~ $1.21 \times 10^{-7}$, which was similar to what obtained before for L1 (Menardo et al. 2019), although with large confidence intervals (Sup. Table 3).

Complementary to the LSD analyses we performed Bayesian molecular clock analyses with Beast 2.5 (Bouckaert et al. 2019). Since Beast analyses are computationally demanding we sub-sampled the

complete L1 dataset to 400 genome sequences, to do this we used two different sub-sampling strategies : 1) random sub-sampling 2) random sub-sampling keeping at least 25 genomes for each year of sampling ("weighted sub-sampling"; see Methods for details). We generated six subsets (three for each sub-sampling strategy), and for each subset we performed a Beast analysis assuming a relaxed clock and an exponential growth prior (Methods).
We found that subsets obtained with the same sub-sampling strategy produced very different results: the highest clock rate estimates was more than 800% larger than the lowest one for the randomly sampled subsets, and more than 100% for the subsets obtained with the weighted sub-sampling strategy (Sup. Table 4). This is a further indication of the lack of a reliable temporal signal for L1.

Since we did not find a reliable temporal signal, we decided to calibrate the phylogenetic tree of L1 with different possible clock rates, and compare the inferred age of MRCAs of some specific clades, for which we have good indications of their earliest possible age from the historical record. More specifically, we considered: 1) clades with a MRCA inferred to have lived in South America, we identified seven such clades composed of at least three strains (Sup. Fig. 5). The phylogeographic analysis inferred these clades to have originated through migration events from the old world (mostly from Africa; Fig. 1, Sup. File 1). It is extremely unlikely that these MTB strains migrated from the old world to South America in pre-columbian times, therefore the MRCA of these clades should not precede the Portuguese discovery of Brazil in the year 1500 AD. 2) We identified one clade endemic to West Africa (Sup. Fig. 5). It consisted of 11 strains sampled in different countries (Ghana, Gambia, Liberia, Mali and Sierra Leone), and it is nested within sublineage L1.1.1, which otherwise, is almost exclusively present in Southeast Asia. The phylogeographic analysis inferred that this clade originated through a direct introduction from Southeast Asia. Again, this is unlikely to have happened before the Portuguese established the maritime route connecting West Africa (and Europe) to Asia, through the circumnavigation of Africa in 1498 AD. Therefore, the MRCA of the West African clade should not precede this time. We calibrated the phylogenetic tree of L1 with different possible clock rate estimates, and tested if the ages of the MRCAs of the Brazilian and West African clade were compatible with the scenarios presented above. To do this we used the dataset used for the phylogeographic analysis, and three different clock rates:
1) $5x10^{-8}$ nucleotide changes per site per year, which is what assumed by O'Neill et al. (2019) in their analysis of L1. This evolutionary rate is very similar to that obtained by the analysis of ancient genomes (Bos et al. 2014), and to some of the Beast results obtained in this study for the complete L1 dataset (although we showed a lack of temporal signal).

2) $1.2 \times 10^{-7}$ nucleotide changes per site per year, this is the clock rate obtained with LSD from the only L1 dataset for which we found a moderate temporal signal in this study (sublineage L1.2.1; Sup. Table 3).
3) $1.4 \times 10^{-7}$nucleotide changes per site per year, this is the clock rate obtained for L1 by Menardo et al. (2019). That dataset showed a strong temporal signal and produced similar results with both LSD and Beast.
We found that the age of the MRCA of the South American clades was posterior to the discovery of Brazil for all three calibration rates. However, the age of the MRCA of the West African clade was earlier than the year 1498 when calibrating the tree with the two lowest clock rates ($5 \times 10^{-8}$ and $1.2 \times 10^{-7}$), while it was posterior when using $1.4 \times 10^{-7}$ as calibration rate (Sup. Fig. 10).
To summarize, we found a lack of temporal structure for L1, the dataset with the best temporal signal (the sublineage L1.1.2; which passed the intermediate DRT) resulted in a clock rate of $1.21 \times 10^{-7}$ nucleotide changes per site per year with LSD. Additionally, clock rates lower than $1.4 \times 10^{-7}$ are not compatible with the hypothesis that the West African clade was introduced directly from Southeast Asia after the first circumnavigation of Africa. In conclusion, although the evidence is weak, these and previous results (Menardo et al. 2019), suggest a clock rate for L1 equal to, or greater than $1.4 \times 10^{-7}$ nucleotide changes per site per year, which means that the MRCA of L1 existed in the 12$^{th}$ century AD or later. We will use this temporal framework for this manuscript, but we emphasize that this is what supported by the available data, and should be regarded as the best possible estimate at this time, additional data, or different methodologies might result in a different scenario.

**Molecular clock analyses of L3**

As for L1, we considered all genome sequences for which the year of isolation was known (827 genome sequences). We used the inferred phylogenetic tree and the years of isolation to time-calibrate the tree with LSD (To et al. 2015), and we performed a date randomization test (DRT) with 100 randomized replicates. We found that the L3 dataset had a good temporal signal (it passed the stringent DRT) and an estimated clock rate of $9.68 \times 10^{-8}$ nucleotide changes per site per year (95% CI: $5.58 \times 10^{-8}$ - $1.21 \times 10^{-7}$; Sup. Fig. 8). These results are similar to previous estimates obtained for L3 (Menardo et al. 2019).
Complementary to the LSD analyses we performed Bayesian molecular clock analyses with Beast 2.5 (Bouckaert et al. 2019), we sub-sampled the L3 dataset to 400 genome sequences, using the same two sub-sampling strategies used for L1 (random sub-sampling and “weighted sub-sampling”). We generated six subsets (three for each sub-sampling strategy), and for each subset we performed two

Beast analyses assuming a relaxed clock and two different tree priors: exponential population growth and the extended Bayesian Skyline Plot (BSP; see Methods for details). In addition, we performed a model selection analysis between these two alternative models, and found that the exponential growth fitted better than the extended BSP for all datasets.

Differently from L1, the results among different datasets obtained with the same sub-sampling strategy were similar, the highest and lowest clock rate estimates between subsets obtained with the same sub-sampling strategy differed at most by 20% (Sup. Table 5). Subsets obtained with the weighted sub-sampling had lower clock rates, while the analysis with the BSP produced higher rates which were closer to those obtained with LSD, however, as mentioned above, the model selection analysis identified the exponential growth prior as best fitting model.

Overall, we found a good temporal signal for L3, nevertheless, the clock rate estimates obtained with different methods were quite variable: the rate obtained with LSD ($9.68x10^{-8}$) was more than double the lowest estimate obtained with Beast ($4.06x10^{-8}$). This is also reflected in the different estimates of the age of the MRCA of L3, which spanned a period between the 2$^{nd}$ and the 13$^{th}$ century AD. If we consider the uncertainty of the estimates obtained with the different analyses, the possible age of the MRCA of L3 was included between the 12$^{th}$ century BC and the 15$^{th}$ century AD (Sup. Table 5).

The large uncertainties and the difference between methods are known limitations of performing molecular clock analyses with MTB datasets, because we calibrate trees with roots that are hundreds, or thousands of years old, with sequences sampled in the last 40 years (Menardo et al. 2019).

As for L1 we will use this range as temporal framework for this manuscript keeping in mind the limitations of such analyses, the large uncertainties, and the assumptions on which these results are based.

**Genome wide scan for positive selection with PAML**

We identified 17 genes inferred to be under positive selection by PAML (see Methods for details), eight in L1, four in L3, and five in both lineages (Sup. Table 9). Seven of these genes were associated with resistance to antibiotics (*RpoB, GyrA, InhA, KatG, EmbB, RpsL,* and *HadA*), and four had unknown function (*Rv2348c, Rv3839, Rv900c, Rv0470A*). The gene *LldD2* was under positive selection in both lineages, this gene was previously identified as under positive selection in MTB (Osorio et al. 2013). We found the effector *EsxM* to be under positive selection in L1, however the positions under positive selection had a large proportion of missing data, and therefore this result could be an artifact caused by misalignment of reads on the reference genome. Finally, we found two genes belonging to the Esx-1 secreting complex (*EccB1* and *EccCa1*), the gene coding for a transmembrane kinase *PknH*, and the gene coding for the secreted antigen *Ald,* under positive selection in L1, but not in L3.

*EsxH* was not among the genes under positive selection in this analysis, although it resulted to be under positive selection when we analyzed a larger set of 500 strains (see main text). This was most likely due to a lack of statistical power: for the genome-wide scan of selection we used datasets with 300 genome sequences, for the analysis of *EsxH* we used a dataset of 500 genome sequences.

**Mutations conferring resistance to antibiotics in L1 and L3**

We considered a set of previously published mutations conferring resistance to 11 different antibiotics (Payne et al. 2019; see Methods). We found that 589 L1 strains (20% of the total), and 516 L3 strains (25% of the total) harbored at least one drug resistance mutation. However, our sample set was based on genome sequences that were publicly available, and it was complemented with additional sequencing. Some of the original studies targeted strains resistant to specific antibiotics, and the prevalence of drug resistant strains in our dataset is larger than what commonly observed in the regions around the Rim of the Indian Ocean (less than 10%; WHO 2019). Therefore, these results should not be taken as a characterization of the global L1 and L3 population, because it is likely that sampling was biased in term of drug resistance.

Nevertheless, we wanted to compare the profile of drug resistance mutations between the two lineages. To mitigate the effect of sampling bias, we did not consider the number of strains harboring a drug resistance mutation, but the number of mutational events along the trees (Methods). We found that the two lineages have similar mutational profiles (Sup. Fig. 12). The most frequently occurring mutation in both lineages was S315T in *KatG,* which represented 20.8% of all mutational events leading to drug resistance for L1, and 22.0% for L3. The main difference between the two lineages was the *InhA* promoter mutation C-15T, which represented 18.0% of all mutational events in L1, but only 5.3% in L3. These results confirm previous findings describing the association of L1 strains with the C-15T mutation at the *InhA* promoter (Fenner et al. 2012).

#### Supplementary Tables

##### Supplementary Table 1

List of samples used in this study, with meta-information.

File: Sup_Table1.xlsx

**Supplementary Table 2.** Results of Mascot analysis for L1 and L3

| **L1** | | |
|---|---|---|
| **Backward migration rate from** | **Median** | **95% HPD** |
| East Africa to Southeast Asia (mainland) | $2.9x10^{-5}$ | $1.3x10^{-8}$ - $1.3x10^{-4}$ |
| East Africa to South Asia | $6.7x10^{-4}$ | $3.4x10^{-4}$ - $1.0x10^{-3}$ |
| East Africa to Southeast Asia (islands) | $9.9x10^{-5}$ | $8.6x10^{-8}$ – $2.6x10^{-4}$ |
| South Asia to Southeast Asia (mainland) | $2.7x10^{-5}$ | $6.8x10^{-9}$ - $1.2x10^{-4}$ |
| South Asia to East Africa | $3.8x10^{-5}$ | $4.9x10^{-9}$ - $1.5x10^{-4}$ |
| South Asia to Southeast Asia (islands) | $1.4x10^{-4}$ | $2.1x10^{-5}$ - $3.2x10^{-4}$ |
| Southeast Asia (mainland) to South Asia | $4.6x10^{-4}$ | $2.1x10^{-4}$ - $7.6x10^{-4}$ |
| Southeast Asia (mainland) to East Africa | $3.3x10^{-5}$ | $3.0x10^{-9}$ - $1.4x10^{-4}$ |
| Southeast Asia (mainland) to Southeast Asia (islands) | $1.6x10^{-4}$ | $2.6x10^{-5}$ - $3.6x10^{-4}$ |
| Southeast Asia (islands) to South Asia | $3.0x10^{-4}$ | $8.8x10^{-5}$ – $5.9x10^{-4}$ |
| Southeast Asia (islands) to East Africa | $3.3x10^{-5}$ | $5.6x10^{-10}$ - $1.5x10^{-4}$ |
| Southeast Asia (islands) to Southeast Asia (mainland) | $1.2x10^{-4}$ | $9.6x10^{-6}$ – $2.9x10^{-4}$ |
| **Effective population size** | **Median** | **95% HPD** |
| East Africa | 5744.3 | 4539.5 – 7163.5 |
| South Asia | 5660.5 | 4621.6 – 6828.1 |
| Southeast Asia (mainland) | 5209.5 | 4106.4 – 6469.2 |
| Southeast Asia (islands) | 4201.5 | 3343.2 – 5199.0 |
| **L3** | | |
| **Backward migration rate from** | **Median** | **95% HPD** |
| East Africa to South Asia | $5.2x10^{-4}$ | $2.6x10^{-4}$ - $8.5x10^{-4}$ |
| South Asia to East Africa | $2.0x10^{-5}$ | $2.3x10^{-11}$ – $8.8x10^{-5}$ |
| **Effective population size** | **Median** | **95% HPD** |
| East Africa | 4224.8 | 3523.9 – 5008.0 |
| South Asia | 10987.9 | 9442.1 – 12724.3 |

**Supplementary Table 3.** Results of LSD analyses for L1

| **Dataset** | **Clock rate** | **Clock rate (95% CI)** | **DRT**[1] |
|---|---|---|---|
| L1 | $3.73 \times 10^{-8}$ | $10^{-10}$ - $6.74 \times 10^{-8}$ | Simple DRT passed |
| L1.1.1 | $10^{-10}$ | $10^{-10}$ - $9.16 \times 10^{-8}$ | DRT failed |
| L1.1.2 | $1.21 \times 10^{-7}$ | $2.53 \times 10^{-8}$ - $1.95 \times 10^{-7}$ | Intermediate DRT passed |
| L1.1.3 | $2.96 \times 10^{-9}$ | $10^{-10}$ - $8.42 \times 10^{-8}$ | DRT failed |
| L1.2.1 | $6.69 \times 10^{-8}$ | $10^{-10}$ - $1.73 \times 10^{-7}$ | DRT failed |
| L1.2.2 | $1.16 \times 10^{-7}$ | $4.84 \times 10^{-8}$ - $1.82 \times 10^{-7}$ | Simple DRT passed |

[1] For the results of the DRT we use the terminology of Menardo et al. (2019): the simple test is passed when the point estimate for the observed data does not overlap with the range of point estimates obtained from the randomized sets. The intermediate test is passed when the point estimate for the observed data does not overlap with the confidence intervals of the estimates obtained from the randomized sets. The stringent test is passed when the confidence interval of the clock rate estimate for the observed data does not overlap with the confidence intervals of the estimates obtained from the randomized sets.

**Supplementary Table 4.** Results of Beast molecular clock analyses for L1

| **Dataset** | **Clock rate** | **Clock rate (95% CI)** |
|---|---|---|
| L1 random 1 | $3.20 \times 10^{-8}$ | $2.51 \times 10^{-9}$ – $8.52 \times 10^{-8}$ |
| L1 random 2 | $6.51 \times 10^{-8}$ | $7.41 \times 10^{-9}$ - $1.25 \times 10^{-7}$ |
| L1 random 3 | $7.87 \times 10^{-9}$ | $2.13 \times 10^{-9}$ - $3.28 \times 10^{-8}$ |
| L1 weighted 1 | $4.39 \times 10^{-8}$ | $1.06 \times 10^{-8}$ – $8.65 \times 10^{-8}$ |
| L1 weighted 2 | $8.83 \times 10^{-8}$ | $2.74 \times 10^{-8}$ - $1.54 \times 10^{-7}$ |
| L1 weighted 3 | $4.19 \times 10^{-8}$ | $7.26 \times 10^{-9}$ - $8.82 \times 10^{-8}$ |

**Supplementary Table 5.** Results of Beast molecular clock analyses for L3

| Dataset | Exponential growth | | Extended bayesian skyline plot | |
|---|---|---|---|---|
| | **Clock rate (95% CI)** | **logML (SD)**[1] | **Clock rate (95% CI)** | **logML (SD)**[1] |
| L3 random 1 | $6.33x10^{-8}$ ($3.27x10^{-8}$ - $1.02x10^{-7}$) | -5450860.3 (28.1) | $8.71x10^{-8}$ ($6.48x10^{-8}$ - $1.09x10^{-7}$) | -5450919.5 (28.2) |
| L3 random 2 | $6.52x10^{-8}$ ($3.78x10^{-8}$ - $9.33x10^{-8}$) | -5458263.6 (27.8) | $8.48x10^{-8}$ ($5.93x10^{-8}$ - $1.08x10^{-7}$) | -5458382.6 (28.2) |
| L3 random 3 | $6.49x10^{-8}$ ($3.82x10^{-8}$ - $9.00x10^{-8}$) | -5468181.6 (27.7) | $8.49x10^{-8}$ ($5.92x10^{-8}$ - $1.05x10^{-7}$) | -5468230.6 (26.8) |
| L3 weighted 1 | $4.85x10^{-8}$ ($3.02x10^{-8}$ - $6.59x10^{-8}$) | -5441674.7 (28.5) | $6.24x10^{-8}$ ($4.61x10^{-8}$ - $8.24x10^{-8}$) | -5441792.0 (28.8) |
| L3 weighted 2 | $4.87x10^{-8}$ ($3.04x10^{-8}$ - $7.16x10^{-8}$) | -5457171.1 (28.4) | $6.42x10^{-8}$ ($4.02x10^{-8}$ - $8.07x10^{-8}$) | -5457315.3 (29.0) |
| L3 weighted 3 | $4.06x10^{-8}$ ($2.16x10^{-8}$ - $6.17x10^{-8}$) | -5456481.0 (28.7) | $5.24x10^{-8}$ ($2.36x10^{-8}$ - $6.83x10^{-8}$) | -5456600.0 (29.3) |

[1] The log marginal likelihood estimated with nested sampling, in parenthesis the standard deviation.

**Supplementary Table 6.** Inferred age of the MRCA of L3

| **Dataset** | Median age of MRCA | 95% CI or HPD of age of MRCA[1] |
|---|---|---|
| L3 complete dataset (LSD) | 1274 AD | 703 AD – 1438 AD |
| L3 random 1 (Beast) | 801 AD | 34 BC – 1371 AD |
| L3 random 2 (Beast) | 866 AD | 218 AD – 1301 AD |
| L3 random 3 (Beast) | 830 AD | 197 AD – 1240 AD |
| L3 weighted 1 (Beast) | 473 AD | 283 BC – 964 AD |
| L3 weighted 2 (Beast) | 500 AD | 259 BC – 1054 AD |
| L3 weighted 3 (Beast) | 184 AD | 1123 BC – 932 AD |

[1] For LSD we report the 95% confidence interval, for Beast we report the interval of the 95% highest posterior density.

**Supplementary Table 7**

Results of the analysis of the diversity of epitopes.

File: Sup_Table7.xlsx

**Supplementary Table 8**

Results of the binding prediction between epitopes and HLA alleles.

File: Sup_Table8.xlsx

**Supplementary Table 9.** Results of PAML. Only genes with Bonferroni corrected p-value < 0.05 are shown

| Gene | M1[1] | M2[2] | LRT[3] | P value | Bonferroni[4] |
|---|---|---|---|---|---|
| **L1** | | | | | |
| *Rv0667 (rpoB)* | -4816.68 | -4750.08 | 133.19 | 1.20E-29 | 4.35E-26 |
| *Rv1266c (pknH)* | -2807.50 | -2793.26 | 28.48 | 6.55E-07 | 2.37E-03 |
| *Rv1484 (inhA)* | -1136.51 | -1124.15 | 24.72 | 4.28E-06 | 1.55E-02 |
| *Rv1792 (esxM)* | -600.45 | -575.25 | 50.41 | 1.13E-11 | 4.11E-08 |
| *Rv1872c (lldD2)* | -1803.30 | -1775.21 | 56.18 | 6.32E-13 | 2.29E-09 |
| *Rv1908c (katG)* | -3119.85 | -3032.51 | 174.67 | 1.18E-38 | 4.27E-35 |
| *Rv2348c* | -508.20 | -495.52 | 25.35 | 3.13E-06 | 1.14E-02 |
| *Rv2780 (ald)* | -1573.18 | -1554.61 | 37.15 | 8.58E-09 | 3.11E-05 |
| *Rv3795 (embB)* | -4700.44 | -4683.43 | 34.02 | 4.11E-08 | 1.49E-04 |
| *Rv3839* | -1059.66 | -1039.36 | 40.60 | 1.52E-09 | 5.52E-06 |
| *Rv3869 (eccB1)* | -2273.40 | -2261.30 | 24.19 | 5.59E-06 | 2.02E-02 |
| *Rv3870 ( eccCa1)* | -3418.58 | -3407.01 | 23.13 | 9.50E-06 | 3.44E-02 |
| *Rv3900c* | -1809.35 | -1776.33 | 66.06 | 4.52E-15 | 1.64E-11 |
| **L3** | | | | | |
| *Rv0006 (gyrA)* | -3650.95 | -3560.72 | 180.46 | 6.52E-40 | 2.36E-36 |
| *Rv0470A* | -619.64 | -608.13 | 23.04 | 9.95E-06 | 3.60E-02 |
| *Rv0635 (hadA)* | -646.02 | -623.03 | 45.98 | 1.03E-10 | 3.75E-07 |
| *Rv0667 (rpoB)* | -5083.81 | -4885.61 | 396.39 | 8.40E-87 | 3.04E-83 |
| *Rv0682 (rpsL)* | -571.69 | -535.97 | 71.46 | 3.05E-16 | 1.10E-12 |
| *Rv1484 (inhA)* | -1032.25 | -1019.41 | 25.67 | 2.66E-06 | 9.65E-03 |
| *Rv1872c (lldD2)* | -1685.78 | -1669.44 | 32.69 | 7.99E-08 | 2.89E-04 |
| *Rv1908c (katG)* | -3442.39 | -3191.79 | 501.18 | 1.48E-109 | 5.35E-106 |
| *Rv3795 (embB)* | -4812.12 | -4661.60 | 301.03 | 4.30E-66 | 1.56E-62 |

[1] The Log-likelihood of the model M1

[2] The Log-likelihood of the model M2

[3] Likelihood ratio test statistic

[4] Bonferroni corrected p-values

**Supplementary Table 10**

Mutations causing resistance to antibiotics in L1 and L3.

File: Sup_Table10.xlsx

#### Supplementary Figures

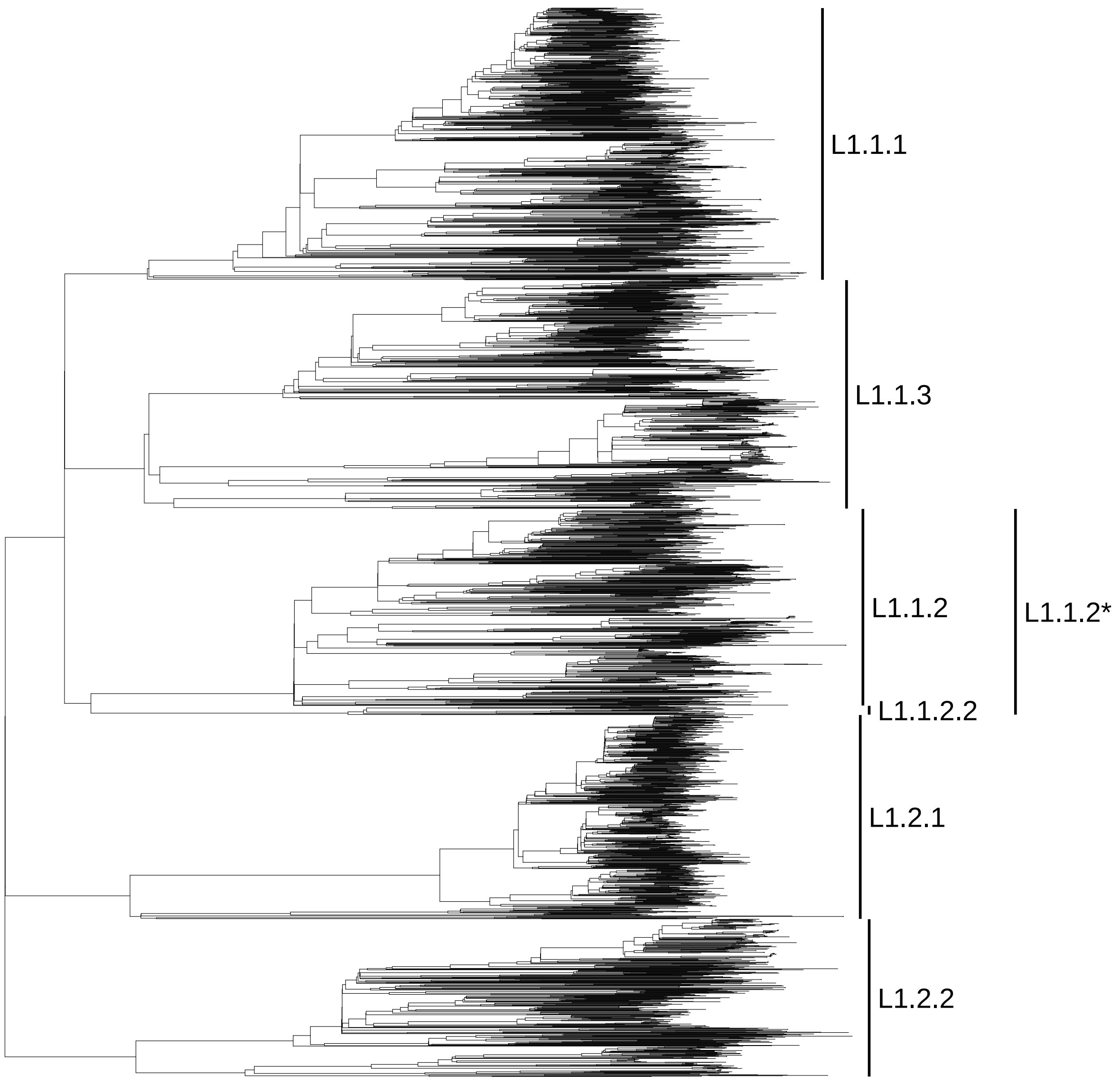


**Supplementary Figure 1**

The phylogenetic tree of the complete L1 dataset (2,938 genome sequences). The sublineages defined by Coll et al. (2014) are labeled. L1.1.2* is the sublineage including L1.1.2 as defined by Coll et al. (2014) and the sublineage L1.1.2.2, which did not fit in the classification of Coll at al. (2014), and was previously identified by Paliitapongarnpim et al. (2018). In this manuscript we refer to L1.1.2* as L1.1.2.

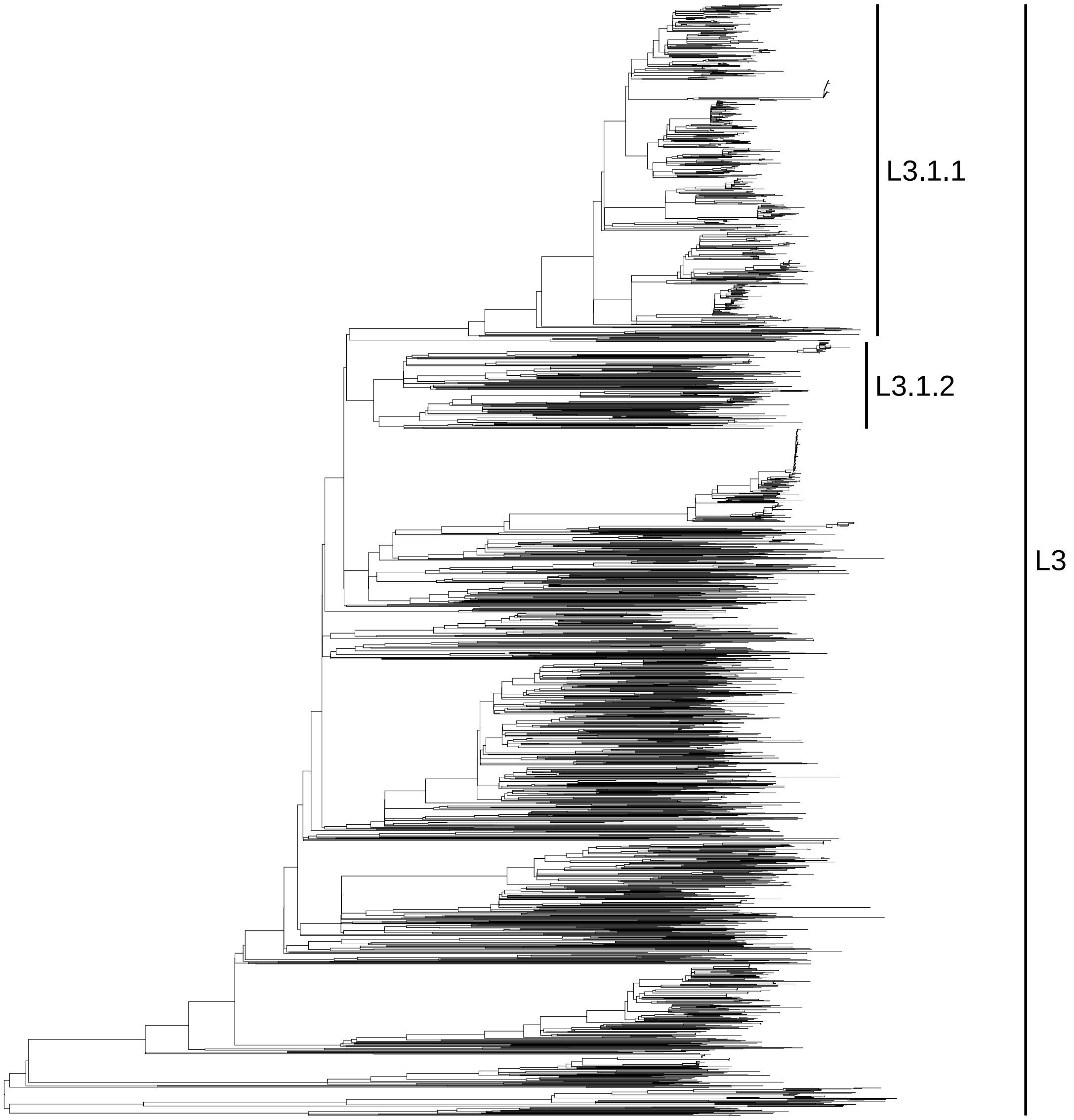


**Supplementary Figure 2**

The phylogenetic tree of the complete L3 dataset (2,308 genome sequences). The sublineages defined by Coll et al. (2014) are labeled.

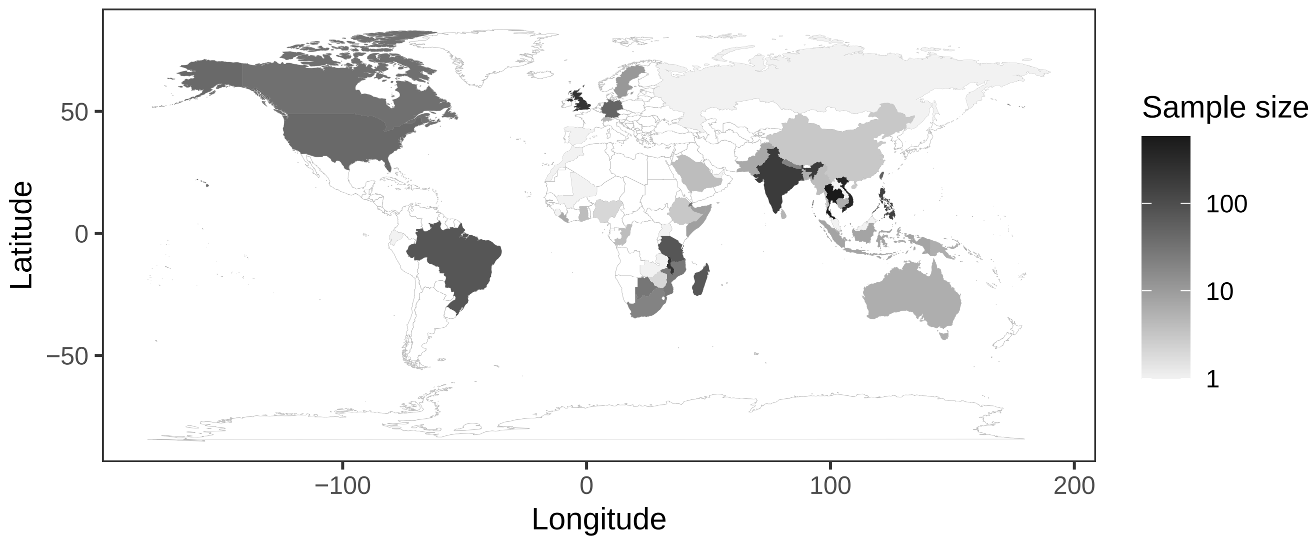


**Supplementary Figure 3**

Heat map showing the country of origin of all L1 sequenced strains for which the information was available (2,451 strains).

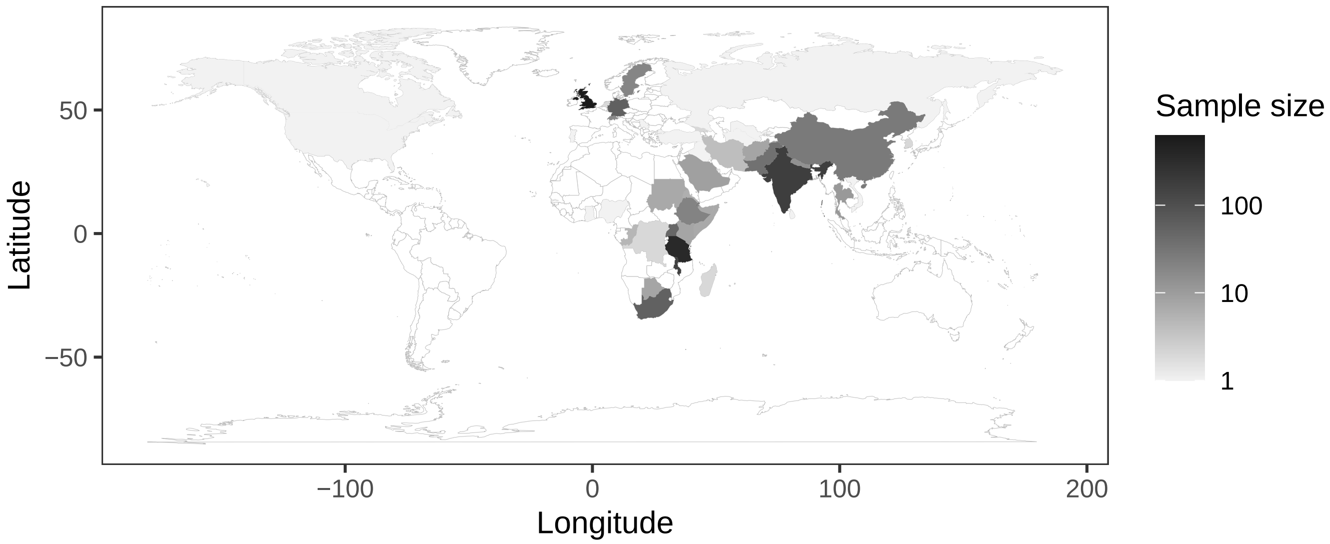


**Supplementary Figure 4**

Heat map showing the country of origin of all L3 sequenced strains for which the information was available (1,748 strains).

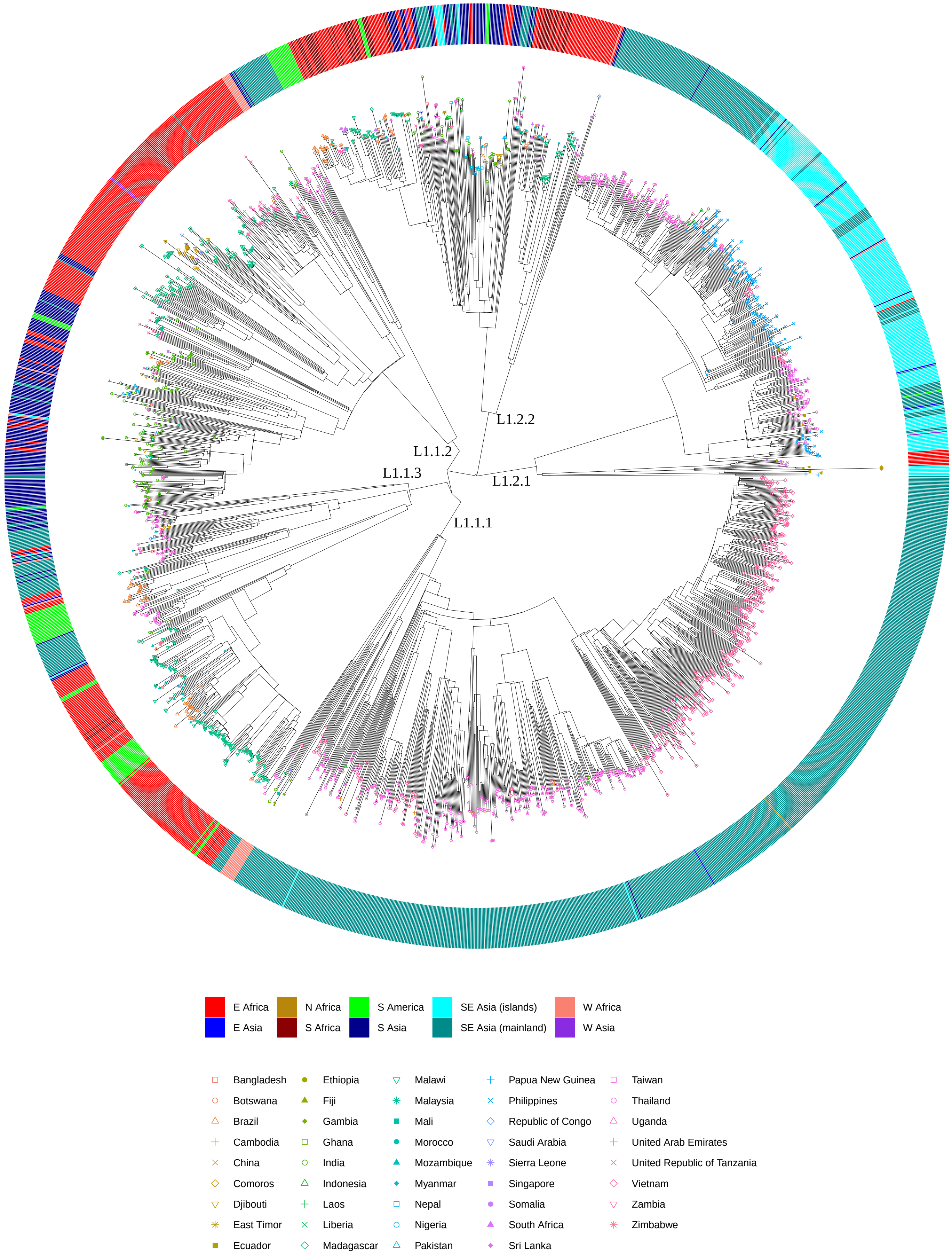


**Supplementary figure 5** (caption in next page)

**Supplementary figure 5**

The phylogenetic tree of the L1 dataset used for the biogeography analysis (2,061 genome sequences). Tips symbols indicate the the country of origin, the heatmap colors indicate the subcontinental geographic region.

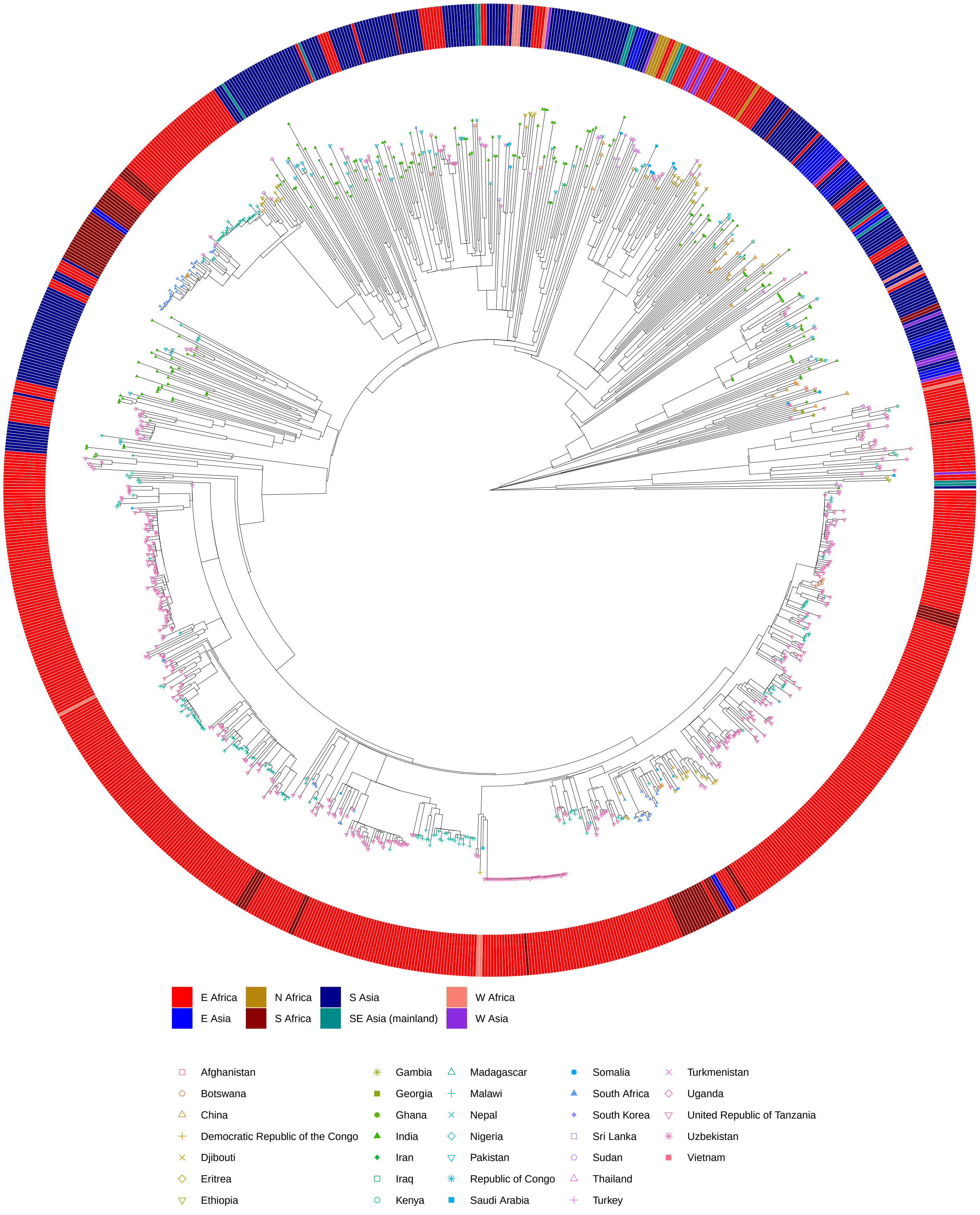


**Supplementary Figure 6** (caption in next page)

**Supplementary Figure 6**

The phylogenetic tree of the L3 dataset used for the biogeography analysis (2,021 genome sequences). Tip symbols indicate the the country of origin, the heatmap colors indicate the subcontinental geographic region.

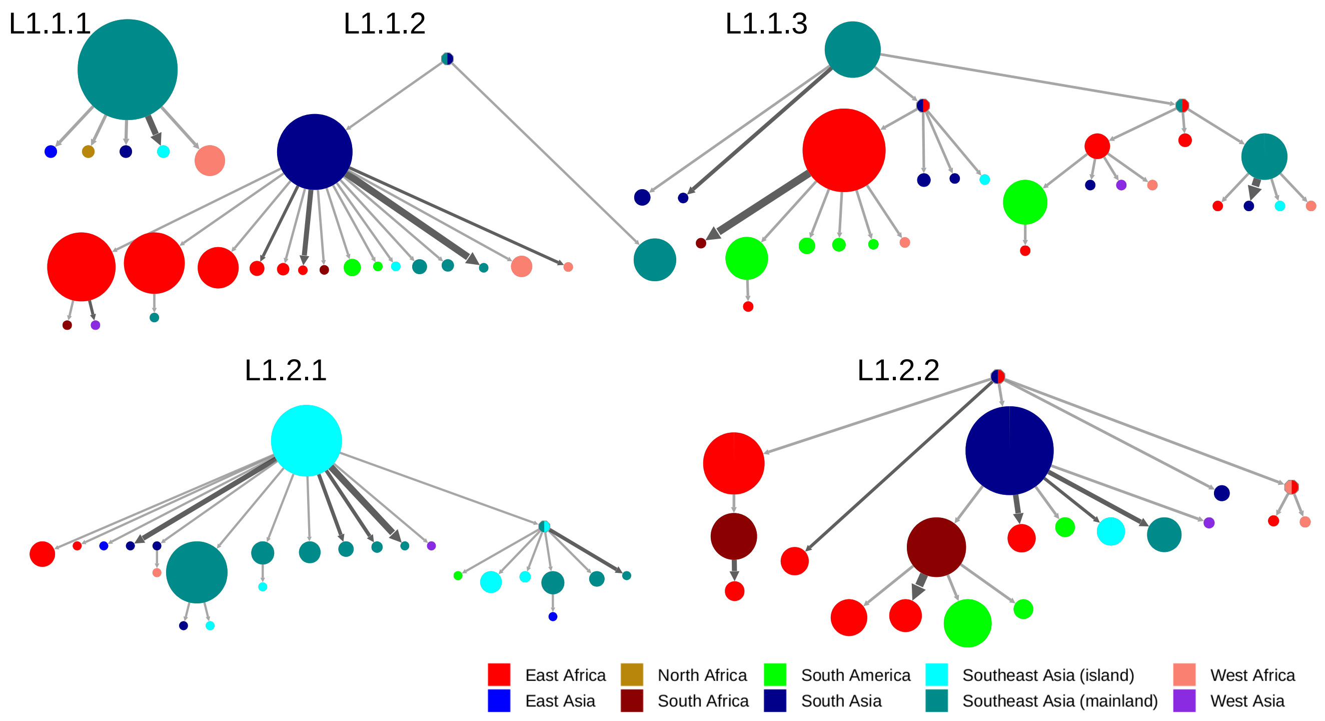


**Supplementary Figure 7**

Results of the PASTML analysis performed on the sublineages of L1 individually.

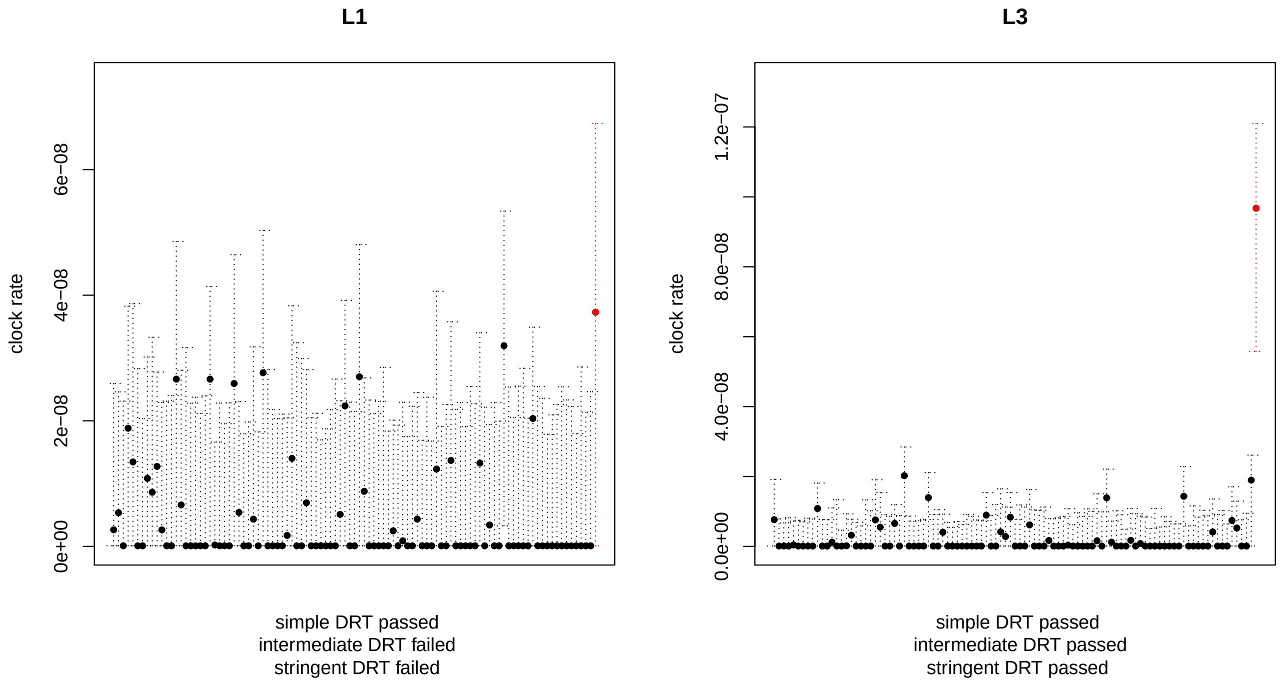


**Supplementary Figure 8**

Results of the DRT with LSD for L1 and L3. The clock rate estimates obtained with observed data are shown in red, the estimates obtained from the randomized replicates are shown in black. The simple test is passed when the point estimate for the observed data does not overlap with the range of point estimates obtained from the randomized sets. The intermediate test is passed when the point estimate for the observed data does not overlap with the confidence intervals of the estimates obtained from the randomized sets. The stringent test is passed when the confidence interval of the clock rate estimate for the observed data does not overlap with the confidence intervals of the estimates obtained from the randomized sets.

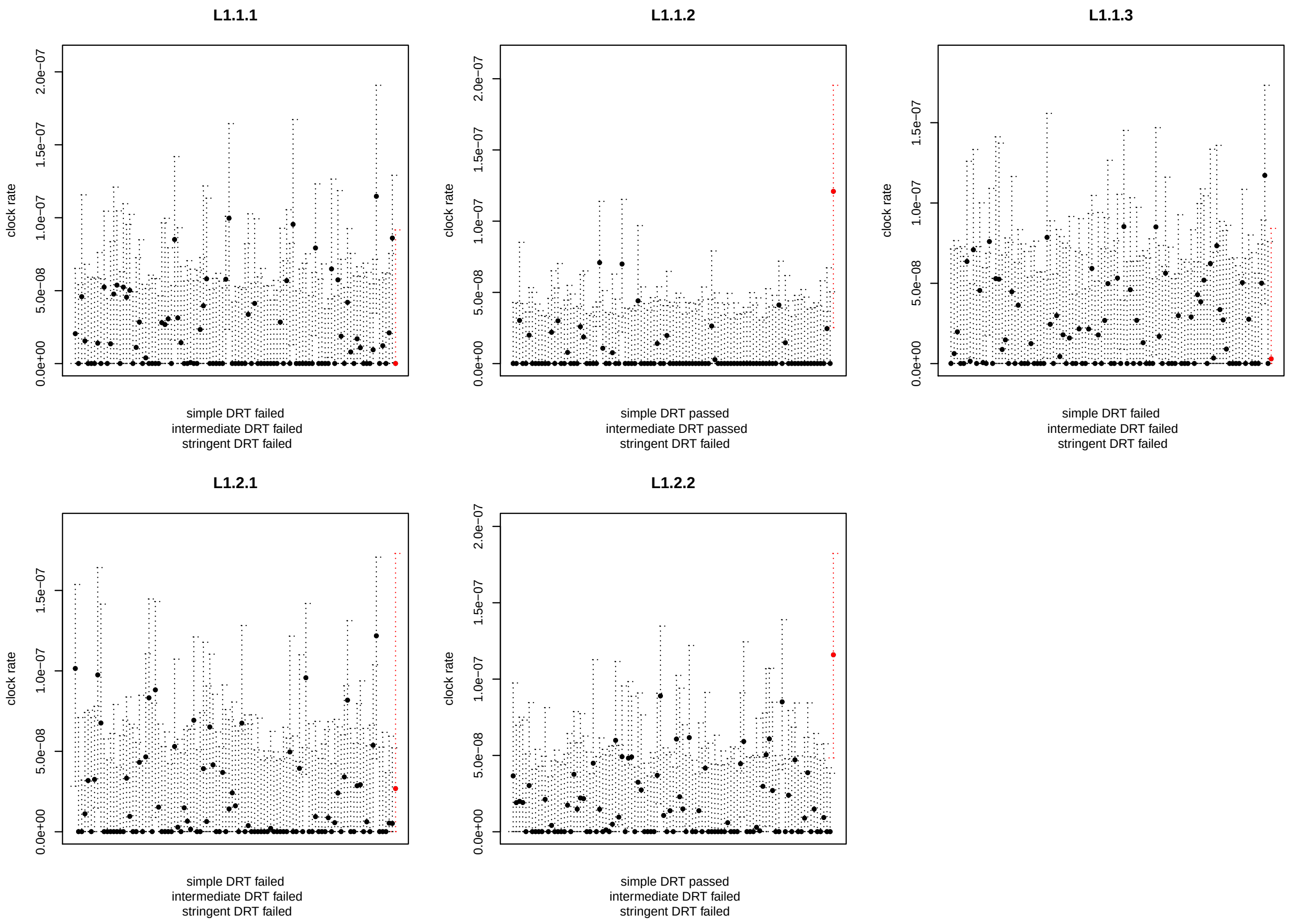


#### Supplementary Figure 9

Results of the DRT with LSD for the sublineages of L1. The clock rate estimates obtained with observed data are shown in red, the estimates obtained from the randomized replicates are shown in black. The simple test is passed when the point estimate for the observed data does not overlap with the range of point estimates obtained from the randomized sets. The intermediate test is passed when the point estimate for the observed data does not overlap with the confidence intervals of the estimates obtained from the randomized sets. The stringent test is passed when the confidence interval of the clock rate estimate for the observed data does not overlap with the confidence intervals of the estimates obtained from the randomized sets.

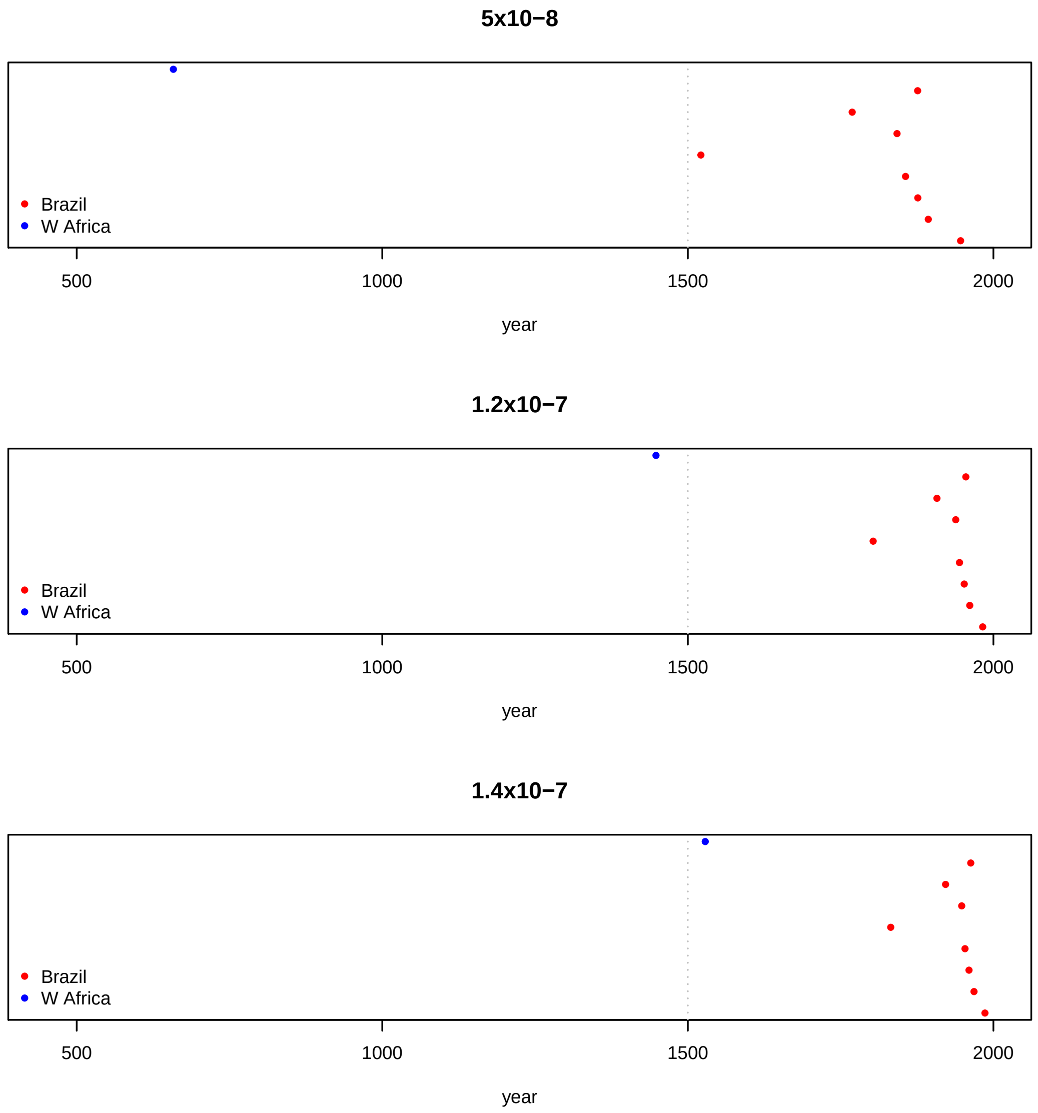


#### Supplementary Figure 10

Estimated age of the most recent common ancestor of the L1 clades endemic to Brazil, and of the West African clade embedded in the sublineage L1.1.1, obtained with three different clock rates.

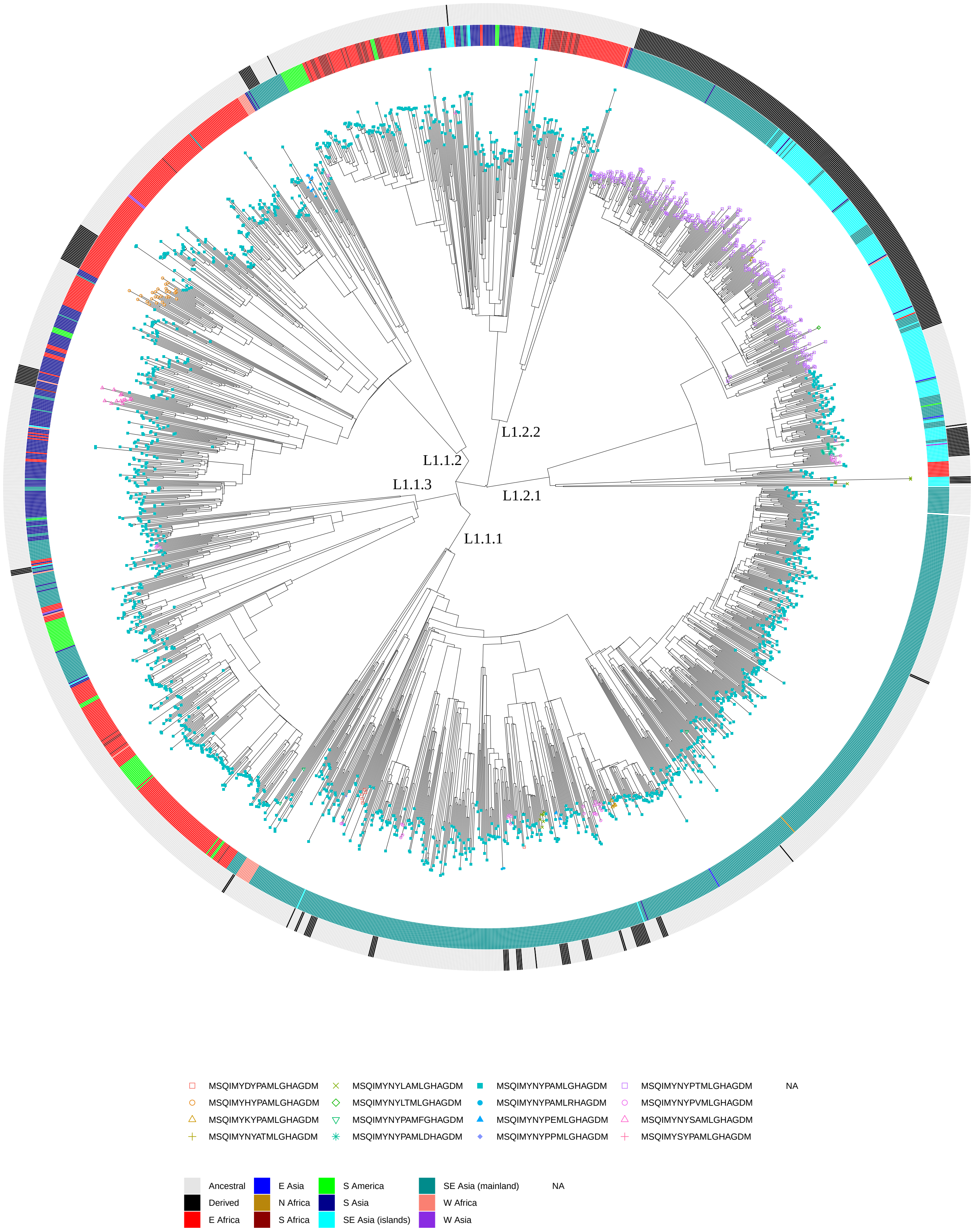


**Supplementary Figure 11** (caption in next page)

**Supplementary Figure 11**

Phylogenetic tree of L1, same dataset used in the biogeography analysis. The tips symbols indicate the haplotype of the N-terminal epitope of EsxH. The colors of the inner ring of the heatmap indicate the geographic region of origin. The colors of the outer ring of the heatmap indicate whether the strain harbor an ancestral or derived haplotype at the N-terminal epitope of EsxH.

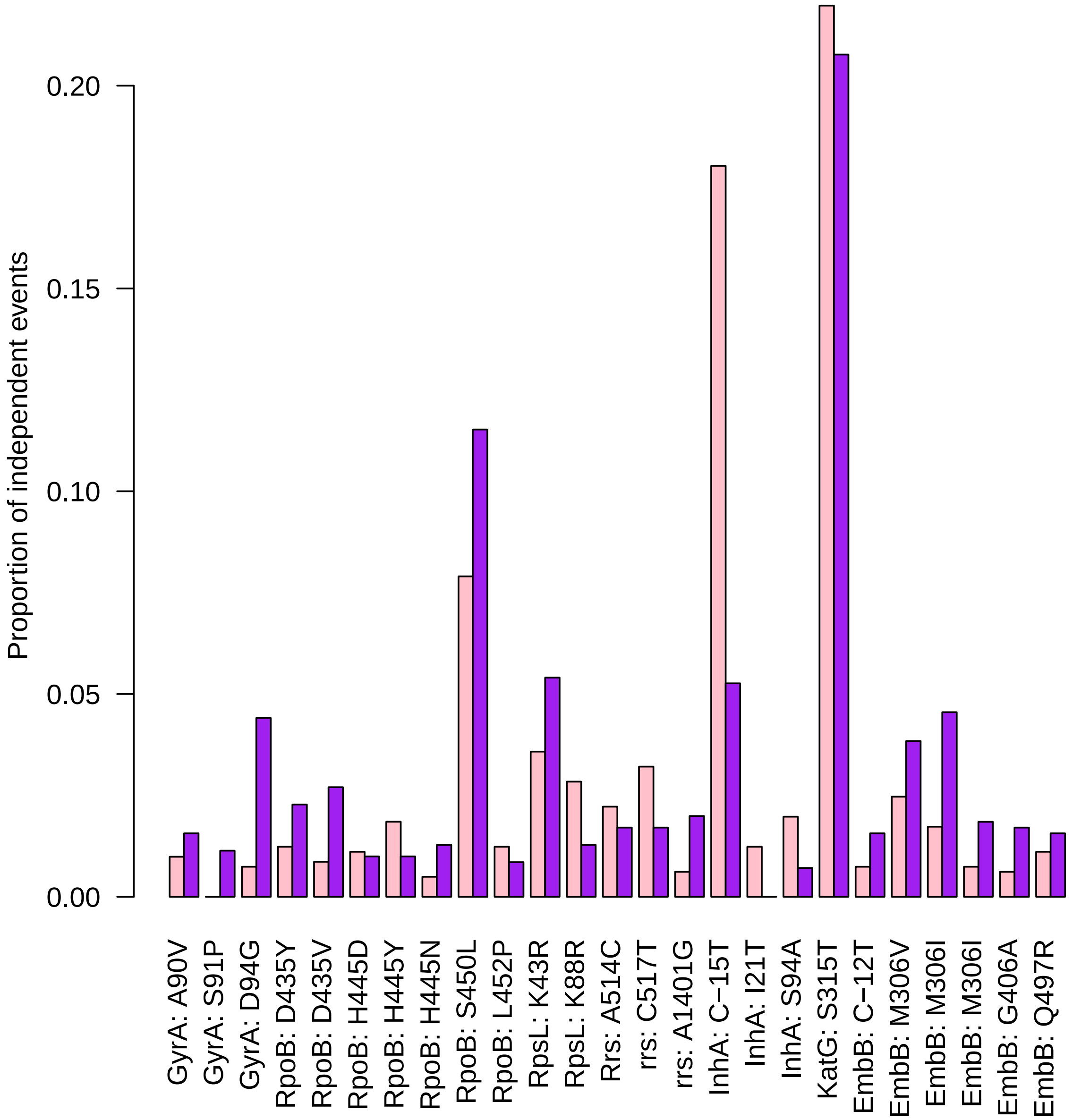


**Supplementary Figure 12**

Mutational profile of drug resistance mutations in L1 and L3. The histogram represent the number of independent mutational events in proportion to the total number of mutational events leading to resistance to any antibiotic. Only mutation accounting for at least 1% of total mutational events in at least one lineage are shown. Mutations in the gene g*yrA* confer resistance to floroquinolones, mutations in the gene R*poB* confer resistance to rifampicin,  mutations in the gene RpsL confer resistance to streptomycin, mutations in the gene *rrs* (ribosomal RNA S16) confer resistance to streptomycin (A514C and C517T) or capreomycin, amikacin and kanamycin (A1401G), mutations in I*nhA* and K*atG* confer resistance to isoniazid, and mutations in EmbB confer resistance to ethambutol.

### References

- Bos, K. I., Harkins, K. M., Herbig, A., Coscolla, M., Weber, N., Comas, I., ... & Campbell, T. J. (2014). Pre-Columbian mycobacterial genomes reveal seals as a source of New World human tuberculosis. *Nature*, *514*(7523), 494-497.
- Bouckaert, R., Vaughan, T. G., Barido-Sottani, J., Duchêne, S., Fourment, M., Gavryushkina, A., ... & Drummond, A. J. (2019). BEAST 2.5: An advanced software platform for Bayesian evolutionary analysis. *PLoS computational biology*, *15*(4), e1006650.
- Coll, F., McNerney, R., Guerra-Assuncao, J. A., Glynn, J. R., Perdigao, J., Viveiros, M., ... & Clark, T. G. (2014). A robust SNP barcode for typing Mycobacterium tuberculosis complex strains. *Nature communications*, *5*(1), 1-5.
- Ishikawa, S. A., Zhukova, A., Iwasaki, W., & Gascuel, O. (2019). A fast likelihood method to reconstruct and visualize ancestral scenarios. *Molecular biology and evolution*, *36*(9), 2069-2085.
- Menardo, F., Duchêne, S., Brites, D., & Gagneux, S. (2019). The molecular clock of Mycobacterium tuberculosis. *PLoS pathogens*, *15*(9), e1008067.
- Müller, N. F., Rasmussen, D., & Stadler, T. (2018). MASCOT: parameter and state inference under the marginal structured coalescent approximation. *Bioinformatics*, *34*(22), 3843-3848.
- O'Neill, M. B., Shockey, A., Zarley, A., Aylward, W., Eldholm, V., Kitchen, A., & Pepperell, C. S. (2019). Lineage specific histories of Mycobacterium tuberculosis dispersal in Africa and Eurasia. *Molecular ecology*, *28*(13), 3241-3256.
- Osório, N. S., Rodrigues, F., Gagneux, S., Pedrosa, J., Pinto-Carbó, M., Castro, A. G., ... & Saraiva, M. (2013). Evidence for diversifying selection in a set of Mycobacterium tuberculosis genes in response to antibiotic-and nonantibiotic-related pressure. *Molecular biology and evolution*, *30*(6), 1326-1336.
- Palittapongarnpim, P., Ajawatanawong, P., Viratyosin, W., Smittipat, N., Disratthakit, A., Mahasirimongkol, S., ... & Kantipong, P. (2018). Evidence for host-bacterial co-evolution via genome sequence analysis of 480 Thai Mycobacterium tuberculosis lineage 1 isolates. *Scientific reports*, *8*(1), 1-14.
- Payne, J. L., Menardo, F., Trauner, A., Borrell, S., Gygli, S. M., Loiseau, C., ... & Hall, A. R. (2019). Transition bias influences the evolution of antibiotic resistance in Mycobacterium tuberculosis. *PLoS biology*, *17*(5), e3000265.
- To, T. H., Jung, M., Lycett, S., & Gascuel, O. (2015). Fast dating using least-squares criteria and algorithms. Systematic biology, 65(1), 82-9.
- World Health Organization. (2019). Global Tuberculosis Report (https://apps.who.int/iris/bitstream/handle/10665/329368/9789241565714-eng.pdf)
